# Does phasic dopamine release cause policy updates?

**DOI:** 10.1101/2022.08.08.502043

**Authors:** Francis Carter, Marie-Pierre Cossette, Ivan Trujillo-Pisanty, Vasilios Pallikaras, Yannick-André Breton, Kent Conover, Jill Caplan, Pavel Solis, Jacques Voisard, Alexandra Yaksich, Peter Shizgal

## Abstract

Phasic dopamine activity is believed to both encode reward-prediction errors (RPEs) and to cause the adaptations that these errors engender. If so, a rat working for optogenetic stimulation of dopamine neurons will repeatedly update its policy and/or action values, thus iteratively increasing its work rate. Here, we challenge this view by demonstrating stable, non-maximal work rates in the face of repeated optogenetic stimulation of midbrain dopamine neurons. Furthermore, we show that rats learn to discriminate between world states distinguished only by their history of dopamine activation. Comparison of these results to reinforcement learning simulations suggests that the induced dopamine transients acted more as rewards than RPEs. However, pursuit of dopaminergic stimulation drifted upwards over a time scale of days and weeks, despite its stability within trials. To reconcile the results with prior findings, we consider multiple roles for dopamine signaling.

## Introduction

An elegant, enormously influential hypothesis on the neural mechanisms of learning^1, 2^ holds that phasic firing of midbrain dopamine neurons encodes reward-prediction errors (RPEs), which optimize action proclivities and value estimates by modifying synaptic weights. This “dopamine-RPE hypothesis” is embedded within a framework developed in the burgeoning field of reinforcement learning^3^ (RL) to interpret key findings from electrophysiological, electrochemical, optogenetic, photometric and behavioral studies. It has been hailed^4^ as “one of the largest successes of computational neuroscience.” Here, we use a free-operant task to conduct a direct test of this hypothesis.

The dopamine-RPE hypothesis was introduced in two seminal papers^1, 2^ that apply the temporal-difference reinforcement-learning (TDRL) algorithm^3, 5^ to foraging behavior in honeybees^1^, to the responses of midbrain dopamine neurons in monkeys performing conditioning tasks^2^, and to rats working for rewarding intracranial stimulation^2^. Most experiments stemming from the dopamine-RPE hypothesis have employed Pavlovian tasks^6–8^ in which the animal has no control over reward delivery. These studies are therefore limited in their ability to determine whether or not phasic dopamine signaling causes behavioral policy updates. Furthermore, the close resemblance between dopamine signals and temporal-difference (TD) RPE signals does not prove that phasic dopamine activity plays the causal role attributed to TD RPEs in updating value estimates and action proclivities^4, 9^. An ideal causal test of the dopamine-RPE hypothesis requires (1) selective and direct manipulation of dopamine activity, (2) an experimental task that gives the subject control over dopamine stimulation and (3) behavioral measurements that reveal changes in action values with high temporal fidelity. Our study achieves these requirements through a multi-step, free-operant task in which rats lever press for selective optogenetic stimulation of their midbrain dopamine neurons. Figure 1 provides a simplified depiction of the hypotheses tested in this paper, regarding the role of dopamine in optical intracranial self-stimulation.

**Fig1:**
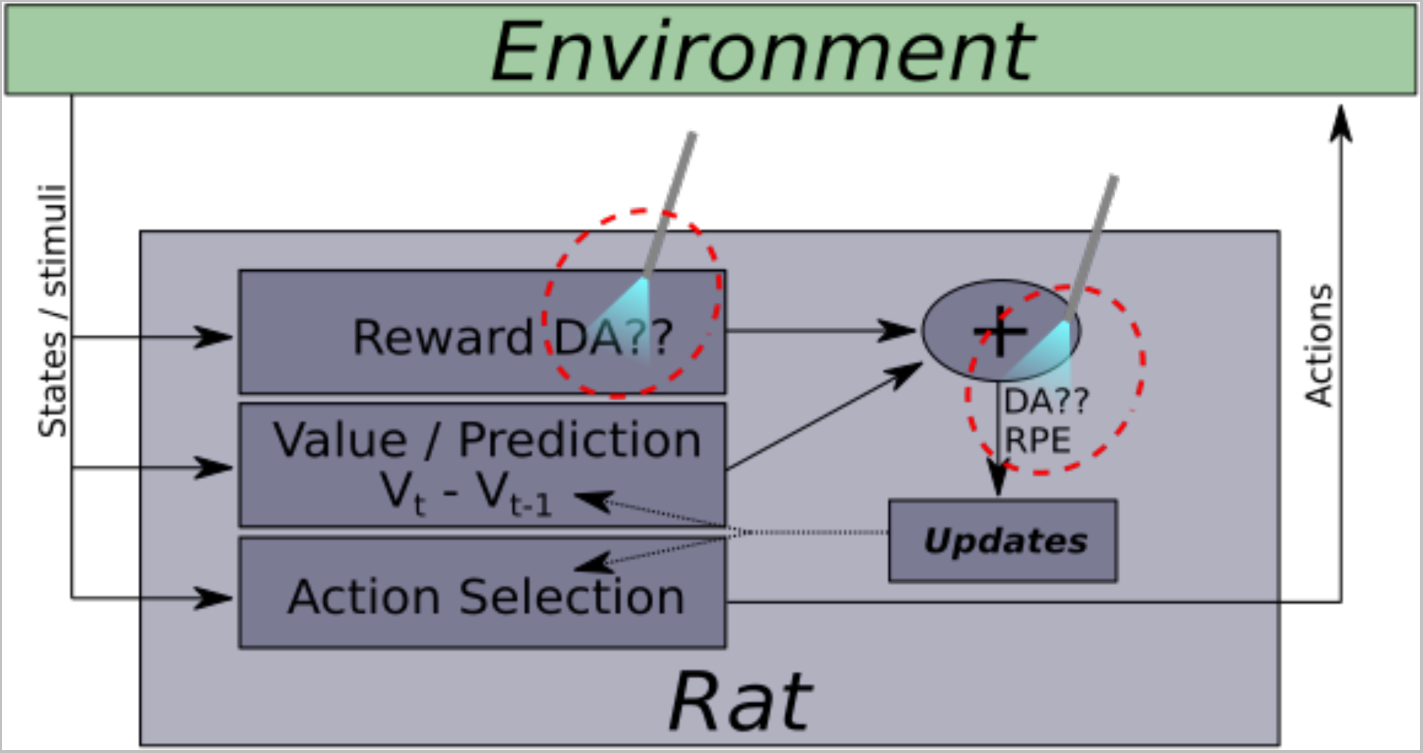
The two competing hypotheses tested here regarding the role of dopamine.

An RPE is commonly interpreted both as an index of how much the value of an outcome exceeds its expectation and as the cause of subsequent behavioral policy changes^10^. In the context of our experiment, this interpretation predicts that optically triggered dopamine transients will act as fictive RPEs. Channelrhodopsin-2 is distributed throughout the cell membrane, including the portion that encloses the axon. Given the small size of the dopamine cell bodies^11^ in comparison to the brain region in which the luminous flux exceeds their firing threshold^12^, the optically triggered dopamine transients cannot be fully nulled by reward predictions provided by inhibitory afferents to the midbrain dopamine neurons (see Figure 1), as we demonstrate by means of voltammetric recordings. In the TDRL simulations of the dopamine-RPE hypothesis reported here, such persistent dopamine transients drive successive, ultimately saturating, increases in value estimates and action proclivities^13^. The stronger the optical stimulation, the higher the expected rate of increase and the lower the number of fictive RPEs required to drive value estimates and action proclivities into saturation. In contrast, if dopamine transients act as fictive rewards, optogenetic stimulation will lead to behavioral updates when the reward is imperfectly predicted, but these updates will diminish progressively as the rat improves its predictions. Thus, reward-seeking behavior will stabilize at non-asymptotic levels.

In summary, if the dopamine-RPE hypothesis is true and the optical stimulation is sufficiently strong to reliably trigger dopamine transients, then the rats will continuously increase their proclivity to press the lever until maximal performance is achieved. Alternatively, if the effect of the dopamine transients is more like that of a reward, then performance for optical stimulation of midbrain dopamine neurons can stabilize at non-asymptotic levels.

The TDRL algorithm is a form of “model-free” learning^2, 14^. It allows an agent to accurately assign values to the various actions available in a given state of the environment. However, this algorithm does not teach the agent how different environmental states are related. Such relations constitute a stored model. In the RL literature, model-free and model-based learning are two poles of a continuum^15^. We show here that learning generated by dopamine activity is not restricted to the model-free pole.

A role for dopamine signaling in model-based learning has been suggested by previous studies ^16–18^. However, these experiments did not isolate dopamine signaling as the sole cause of learning. In contrast, in the present study, dopamine transients provide the sole signal that distinguishes world states.

This experiment poses two broad questions. 1) Does phasic dopamine signaling act more like an RPE or a reward in rats working for optical stimulation of midbrain dopamine neurons? 2) Is the learning induced by dopamine transients confined to model-free RL? To answer these questions, we trained rats to work for optical stimulation of midbrain dopamine neurons. The rats held down a lever in order to trigger the stimulation. Experimental sessions consisted of cycling triads of trials (Figure 2b). During the leading trial of each triad, the strength (pulse frequency) of the optogenetic stimulation was set to a value (High) that elicited maximal performance, whereas during the trailing trial, it was set to a value (Low) that elicited minimal performance. The stimulation strength on offer during the central (”test”) trial of each triad was also constant within a trial but was selected randomly from a set of values that elicited either maximum, intermediate, or minimal performance; the maximum and minimum values were the strengths used in the leading and trailing trials, respectively. In Experiment 1, different levers distinguished test trials from leading or trailing trials. In Experiment 2, no external cues were provided to help the rats discriminate between trial types, making the stimulation-induced dopamine transients the sole signal available to drive learning of the trial structure. A visual cue (illumination of a light over the lever) confirmed that the lever was depressed. Figure 2b describes the sequence of trials, and Figure S1 provides a more detailed illustration of the testing paradigms employed in Experiments 1 and 2.

**Fig2:**
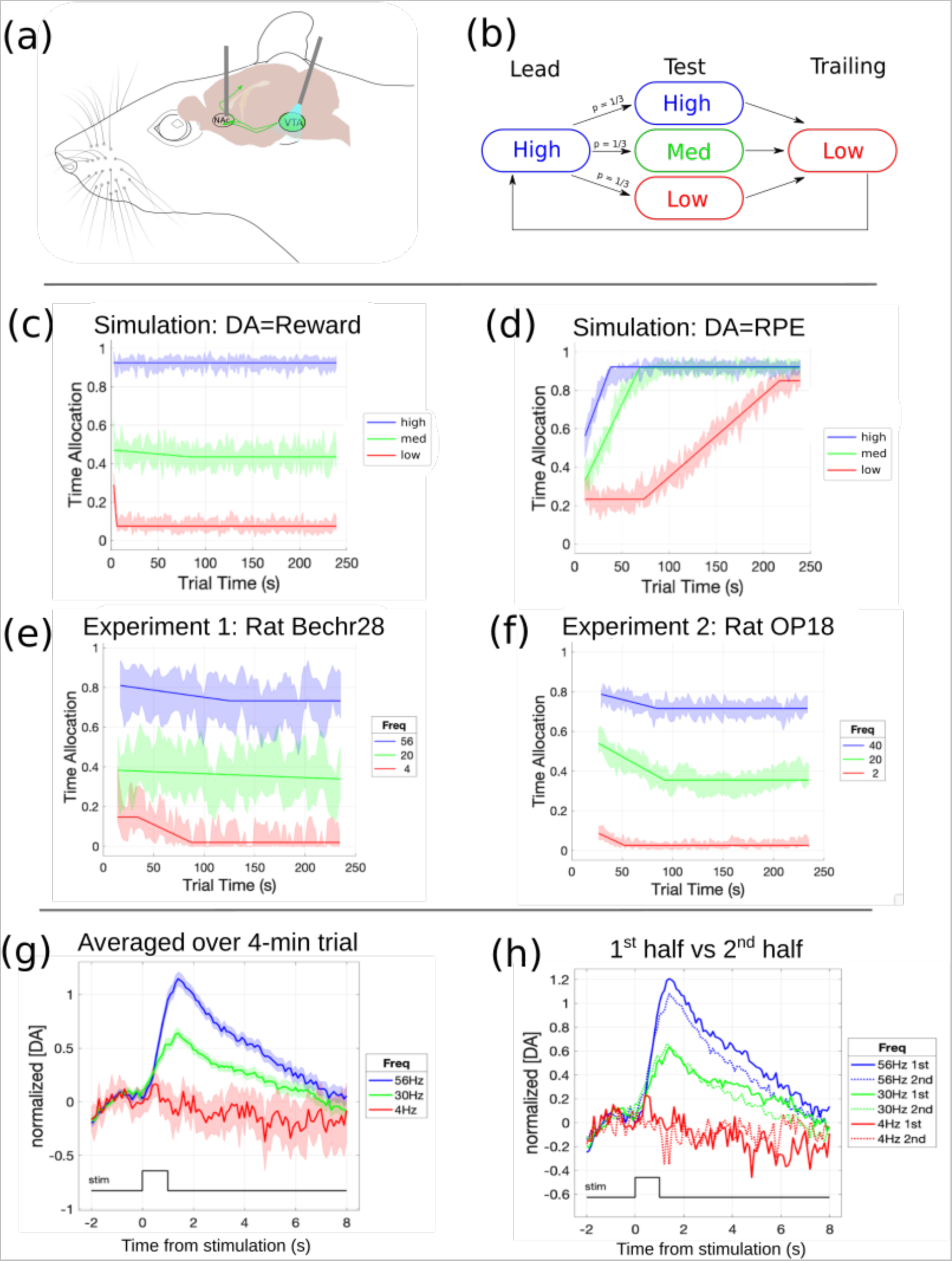
Experimental setup, comparison of simulated and empirical results, and fast-scan cyclic voltammetry data. **a**: implanted optical fiber for optogenetic stimulation of midbrain dopamine neurons and fast-scan cyclic voltammetry electrode for recording of dopamine transients in the nucleus accumbens (image of rat head and brain from https://neuroscience-graphicdesign.com/portfolio/laboratory-rats/); **b**: trial structure. High, Med, and Low refer to high, medium and low pulse frequencies, respectively; **c**: simulated time allocation to pursuit of High, Med, and Low strength trains, assuming that the optically triggered dopamine transients acted as rewards; **d**: simulated time allocation to pursuit of High, Med, and Low strength trains, assuming that the optically triggered dopamine transients acted as RPEs; **e**: observed time allocation to pursuit by rat Bechr28 of High, Med, and Low pulse-frequency trains in test trials (Experiment 1); **f**: observed time allocation to pursuit by rat OP18 of High, Med, and Low pulse-frequency trains in test trials (Experiment 2); **g**: normalized dopamine concentrations from fast-scan cyclic voltammograms obtained in response to High, Med, and Low pulse-frequency trains; **h**: same data as in **g**, but split between the first and second half of the trials.

As shown by our simulations, stable performance at intermediate levels is incompatible with model-free TDRL, under the dopamine-RPE hypothesis. However, we observed such performance consistently.

On test trials, the rats could not know which stimulation strength was on offer until they had earned at least one reward. In contrast to the slow, incremental updating typical of model-free learning, we show that test-trial performance was adjusted very rapidly, sometimes after only a single reward had been obtained. In Experiment 2, the history of stimulation-induced dopamine activity provided the only basis for guessing the identity of the current trial type prior to delivery of the first reward on the trial. All rats showed differential responding across trial types prior to delivery of the first reward, and some guessed the trial type reliably.

## Results

### Phasic dopamine activity does not cause policy updates

To address the causal role of dopamine transients in updating the stored values underlying reward seeking, we determined how time was allocated to self-stimulation in test trials, across high, medium (Med), and low stimulation strengths (Figure 2e,f; S2,S3). The proportion of time that the lever was depressed is shown in successive 2-s time blocks. Lines show best-fitting piecewise-linear functions, and error bands show the surrounding 95% confidence intervals. To ensure that there are roughly comparable numbers of observations in each 2-s time block, the plots for each stimulation strength begin at the time block when responding had commenced on 75% of the trials. Due to a software bug, some trials terminated prematurely by up to 4s (2 time blocks). Thus, the ends of the plots are truncated at the third to last time block.

As expected, the rats worked maximally for the high-strength stimulation, negligibly for the low-strength stimulation, and typically at intermediate levels for the medium-strength stimulation. Many more test sessions were run in Experiment 2 than in Experiment 1, and the width of the error bands reflect this.

Critically, in all rats, the trajectory of responding did not accelerate upwards (Figure 2e,f; S2,S3), contrary to the predictions of the TD-RL simulations of the dopamine-RPE hypothesis (Figure 2d). Altering the simulation such that dopamine acts as a reward led to better predictions of the rats’ actual behavior (Figure 2c).

The roughly stable or declining time allocation over the course of the trial (Figures 2e,f, S2,S3) was not an artifact of averaging across trials. As shown in Figures S4,S5, Rats distributed their time between work (holding down the lever) and leisure (e.g., grooming, resting, exploring). The duration of work bouts typically increased, and the duration of leisure bouts typically decreased, as a function of increasing stimulation strength.

Fast-scan cyclic voltammetry (FSCV)^19^ was used to monitor stimulation-evoked dopamine release in the nucleus-accumbens terminal field in a rat performing the triadic-trial task (Figure 2g,h). These results parallel the behavioral data in Figure 2e,f. The magnitude of the dopamine transient evoked by the optical stimulation increased as a function of stimulation strength and remained roughly stable over the entire course of the 4-min trial. Note that peak dopamine release evoked by the medium-strength train was of similar amplitude during the first and second halves of the trial. A train-by-train plot of the peak dopamine concentrations is shown in Figure S13.

Video S1 shows stimulation-induced dopamine release and lever-pressing behavior in the triadic-trial task.

### Dopaminergic stimulation sufficed to support trial-cycle tracking and prediction

To evaluate the presence or absence of knowledge regarding the triadic organization of trials, we compared the distribution of initial response latencies across leading, test, and trailing trials (Figure 3, S6, S7). We reasoned that learning and subsequently tracking the trial sequence over the course of the experiment would motivate a rat to respond fastest at the start of a leading (high-stimulation-strength) trial, slowest at the start of a trailing (low-stimulation-strength) trial and with an intermediate latency at the start of test trials (when either high-, medium-, or low-strength stimulation would be on offer). Not surprisingly, the proportion of subjects that learned the trial sequence correctly was higher in Experiment 1 (4 of 5 rats), when test trials were signaled by extension of a lever used uniquely on those trials (Figures S6), than in Experiment 2 (2-3 / 6 rats), when the same lever was extended on all trials, and no stimuli were available to indicate the identify the trial type (Figures 3b,f;,S7). Nonetheless, all rats in Experiment 2 responded fastest at the start of lead trials. The differential initial latencies shown by the rats demonstrate that dopamine signaling was sufficient to track the trial structure, either partially or fully.

**Fig3:**
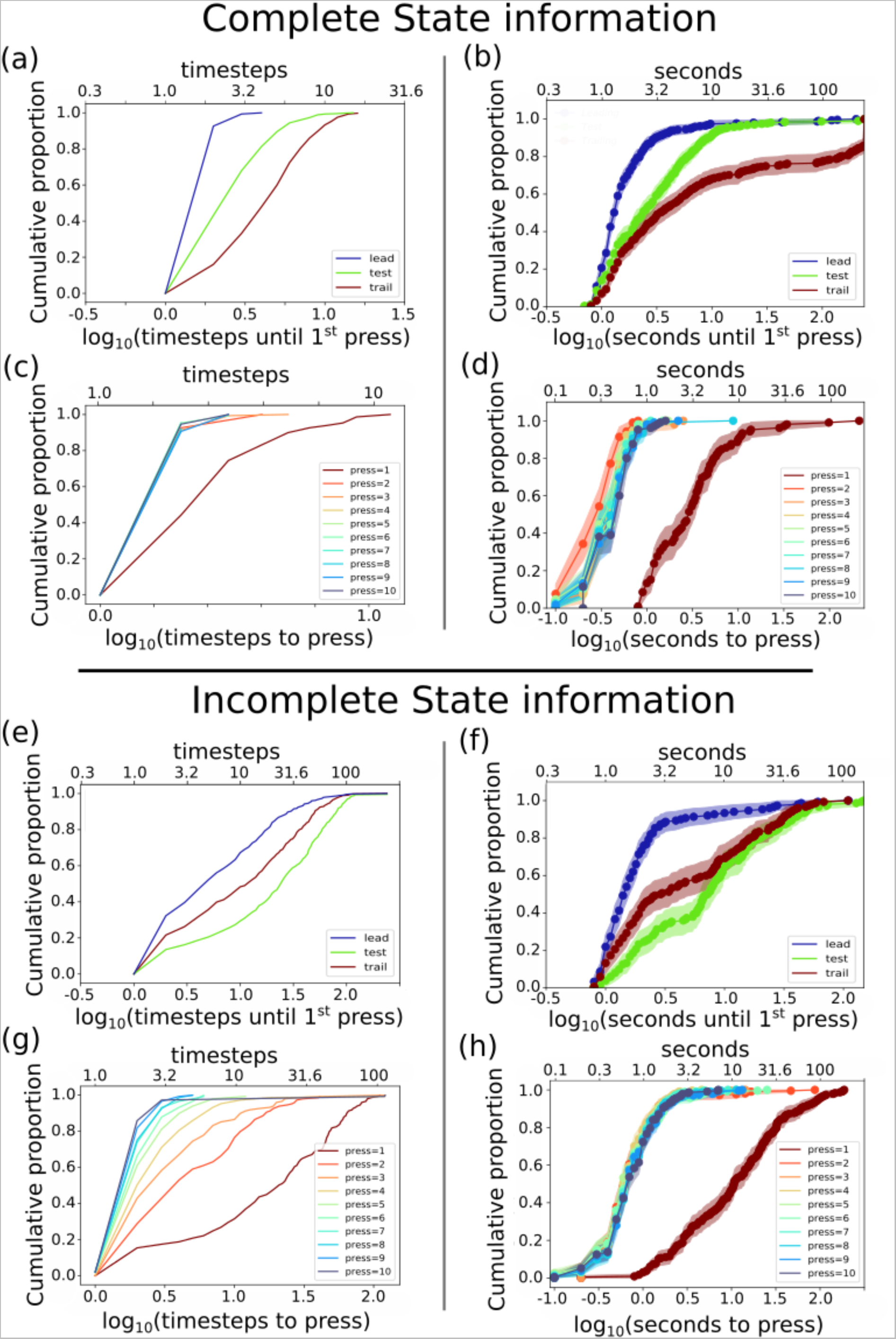
model formation: **a:** cumulative distribution of latencies until first lever press in a simulation with an agent that has perfect state information. **b:** cumulative distribution of latencies until first press in rat OP13. The increasing latencies from lead to test to trailing trials show that the rat understood and tracked the triadic-trial structure **c:** cumulative distribution of latencies between presses in a simulation with an agent that has perfect state information. **d:** cumulative distribution of latencies between presses in high-strength test trials for rat OP13. **e:** cumulative distribution of latencies until first press in a simulation with an agent that has incomplete state information. **f:** cumulative distribution of latencies until first press in rat OP18. The shorter latencies in trailing trials than in test trials show that the rat had an imperfect understanding of the triadic-trial paradigm **g:** cumulative distribution of latencies between presses in a simulation with an agent that has incomplete state information. **h:** cumulative distribution of latencies between presses in high-strength test trials for rat OP18.

Following and extending earlier work by Breton^20^, Ahilan and Dayan analyzed the performance of rats working for rewarding electrical brain stimulation in a fuller version of the triadic-trial paradigm^21^. These analyses show that when the stimulation strength and/or opportunity cost of the stimulation on a test trial differed markedly from those on leading and trailing trials, the rats tracked the trial sequence well. In contrast, the rats appeared to confuse test trials with either leading or trailing trials when the conditions were similar. This led them to initiate responding with short latencies on the subsequent trailing trial. This tendency to quickly initiate responding on a trailing trial following a low-stimulation-strength test trial is also seen in the data from the two rats in Experiment 2 that most accurately learned the trial sequence, ELOP21 and OP13 (Figure S8). Initial response latencies for the remaining rats were also lowest on leading trials, but tended to be shorter than those on test trials in the cases of rats ELOP18, ELOP22, and OP18 (Figure S7). Varying the state representation and state transitions(Figure 3e,g) allows the simulations to replicate the observed difference in the ability of the rats to learn and track the trial sequence. (See methods section for details).

Figure 3d extends the analysis of response latencies to the first 10 bouts of rewarded lever depression on test trials. This extended analysis reveals the rate at which behavior adjusted following resolution of the initial uncertainty about the strength of the stimulation on offer and sheds light on whether the rats had learned that stimulation strength remains constant throughout a given trial.

As shown in Figure 3d, rat OP13 (Experiment 2) showed extremely rapid updating on high-stimulation-strength test trials. At the start of a test trial, the rat could not know which of the three possible stimulation strengths would be available. This uncertainty is reflected in the position of the cumulative distribution function for the first bout, which lies well to the right of the subsequent ones, with a mean latency of ∼3 s. After the high stimulation strength was revealed by delivery of the first stimulation train, the cumulative distribution function jumped leftwards; the mean latency to press the lever declined roughly tenfold and remained roughly stable thereafter.

### Valuation of dopamine stimulation grew slowly over extended testing

In Experiment 2, rats were tested repeatedly in the triadic-trial paradigm for an extended period spanning many weeks. Over the course of repeated testing, five of the six rats showed a reliable increase in the time spent working for the medium-strength stimulation, and the sixth (ELOP18) showed a trend in the same direction (Figures 4a, S10). We offer two, possibly complementary, hypotheses about the reason for this upwards drift. First, we add some pertinent data.

**Figure 4:**
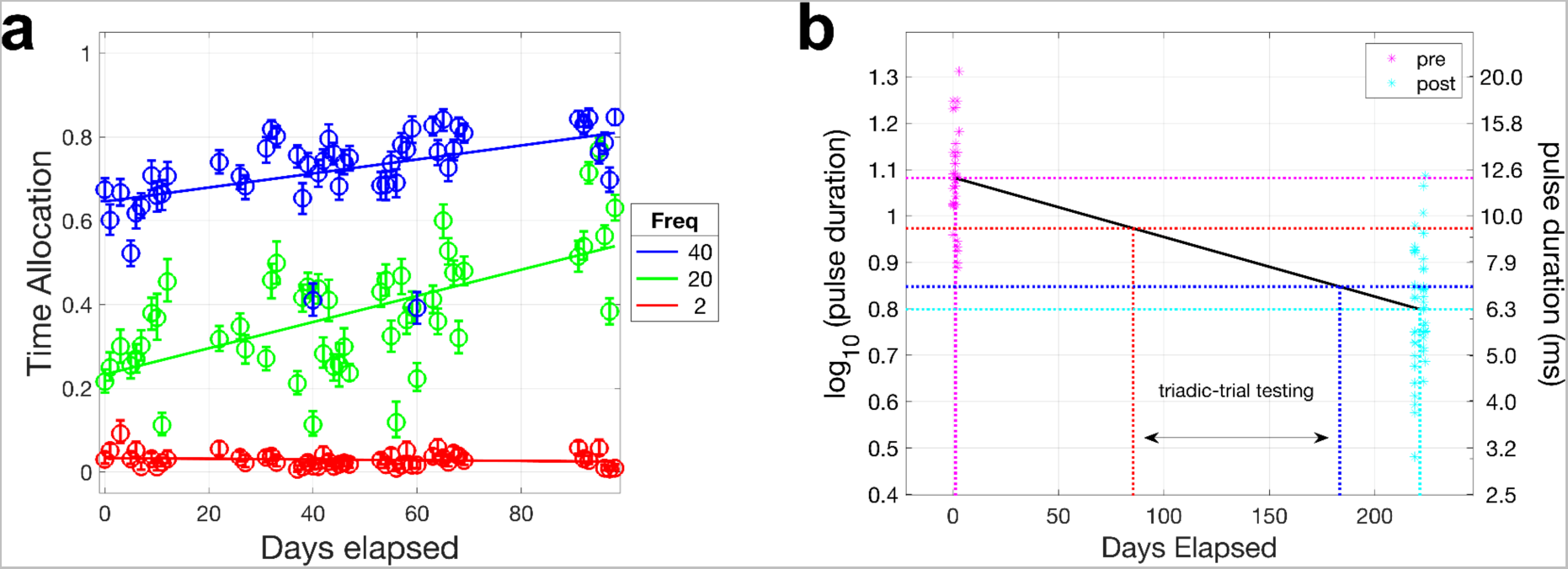
Drift of time allocation over the course of repeated testing in rat OP18. **a:** Time spent working as a function of repeated testing for high-, medium- and low-strength stimulation trains. **b.** Position-parameter values of sigmoidal functions fitted to data from pulse-duration sweeps performed before and after the triadic-trial experiment. Lower pulse durations indicate higher rewarding effectiveness. Position-parameter values at the start and end of triadic-trial testing were interpolated on the basis of the linear trend between the two pulse-duration sweeps.

Prior to and following testing in the triadic-trial paradigm, data were obtained from four Experiment-2 subjects showing how time allocation varies as a function of the duration of the optical pulses delivered to the ChR2-expressing midbrain dopamine neurons (Figure S11). The location of the time-allocation versus pulse-duration curves along the pulse-duration axis reflects the behavioral effectiveness of the stimulation^22^: the shorter the pulse duration required to achieve a given level of time allocation, the more “bang” for the optical “buck.” In all four cases, the curves shifted leftwards (towards shorter durations) between the pre- and post-triadic-trial testing, indicating that the optical stimulation had become more effective.

The time-allocation versus pulse-duration data from two subjects were obtained at the same pulse frequency (20 Hz) that was used in triadic-trial testing as the medium stimulation strength. Sigmoidal curves were fitted to both of these datasets. As explained in detail in the supporting information, the parameters of these curves were used to predict the drift in time allocation for the medium stimulation strength in Experiment 2. Figure S11 shows that this prediction accounts for 64% of the observed drift in Figure 4a at the medium stimulation strength (green line). In the case of rat OP20, the same calculation accounts for 36% of the observed drift. These data show that the increasing time allocation over the course of testing was not specific to the triadic-trial task.

## Discussion

The RPE is the engine of systematic change in reinforcement-learning theories. Following a shift in reward magnitude, the RPE adjusts expectations and action weights accordingly. As long as RPEs are generated consistently, expectations and behavior should undergo continued adjustment until weights are driven into saturation. Expectations and action weights stabilize only when the RPE disappears.

According to the dopamine-RPE hypothesis, burst firing in midbrain dopamine neurons encodes RPEs and causes the learning and behavioral changes they engender. The results reported here challenge this causal proposition. We show that contrary to the dopamine-RPE hypothesis, rats worked for many minutes at stable, non-maximal levels for optical activation of midbrain dopamine neurons. The rapid adjustments following receipt of the initial stimulation trains on test trials show that the behavior of the rats could and did change abruptly. Thus, the stability of test-trial performance after the initial adjustment cannot readily be attributed to high behavioral inertia.

The observed behavioral stability was mirrored in FSCV recordings of dopamine release during performance in the triadic-trial paradigm. Over the course of test trials during which a rat earned 49-61 trains of optical pulses, stable, peak dopamine concentrations were observed in the nucleus-accumbens terminal field in response to the medium stimulation strength, and only a minor decline was observed in response to the high stimulation strength.

We used the TDRL algorithm^2, 3, 5^ to simulate the behavior expected on the basis of the dopamine-RPE hypothesis in response to stable stimulation-induced release of dopamine. This widely-used algorithm provides a simple implementation of RPE-based learning. It predicts that when an action repeatedly and consistently generates an RPE, the probability of selecting that action will grow progressively until an asymptotic level is achieved. This prediction stands in sharp contrast to the observed stability of responding in the present experiments. Altering the simulation so that dopamine-release events mimic primary rewards, rather than RPEs, yields results more congruent with the behavioral findings.

### Model-free versus model-based learning

Different ways in which the fruits of prior learning can control ongoing behavior have been arrayed along a continuum stretching from a simple “model-free” pole to a more elaborate and computationally demanding “model-based” pole^15^. The TDRL algorithm lies at the model-free pole. At the opposite pole, the agent stores a representation that allows it to explore prospectively^15^. The data in Figures 3b,f,S7 suggest a form of learning positioned beyond the model-free pole. Although the paradigm cannot provide evidence of prospective exploration, the initial latencies do show that the rat could predict “how the environment will respond to its actions”^3^ on the basis of dopaminergic activation produced on previous trials.

In Experiment 2, no external cues were provided to distinguish leading, test, and trailing trials. Nonetheless, most initial response latencies on leading trials were much shorter than on the other two trial types. Thus, all of these rats were able to track the phase of the triadic-trial cycle with at least partial accuracy, predicting correctly that a leading trial had begun before they had earned a reward on that trial. There is strong evidence that two rats {ELOP21, OP13} successfully tracked all three phases and weaker evidence that a third (OP20) did so as well.

The sole feature that distinguishes the trial types in Experiment 2 is the optically induced firing of midbrain dopamine neurons. Thus, the stored history of this transient neural signal was the basis for the rats’ predictions. This shows that in addition to any role in the *mechanism* of learning, the transient firing of dopamine neurons can be mapped into enduring *content* within learned representations.

A distinction can be made between “strong” and “weak” forms of the dopamine-RPE hypothesis. The strong form holds that dopamine RPEs directly update action values, whereas a weak form could implicate dopamine-mediated RPE signals in other processes such as Pavlovian, but not instrumental, learning. We argue against the strong form, and we also note a feature of our experiment that seems inconsistent with the abovementioned weak form. A cue (illumination of the lever light) accompanied lever pressing. If the optically induced activation of midbrain dopamine neurons acted as an RPE that adjusted Pavolvian associations, wouldn’t Pavolvian-instrumental transfer^23^ be expected to invigorate the lever-pressing behavior, thus increasing hold times and time allocation? Instead, time-allocation remained stable or mildly decreasing over many minutes.

### Compensatory inhibition of unstimulated dopamine neurons?

The unilateral optical stimulation cannot reach all midbrain dopamine cells. Could inhibition of unstimulated dopamine neurons outside the effective stimulation field have compensated for the repeated activation of the ChR2-expressing subpopulation near the tip of the optical fiber? On this view, unstimulated neurons would come to be inhibited by the reward predictions, ultimately holding constant the net activity of the dopaminergic population and eliminating the RPE.

We regard this as unlikely. In an fMRI study, blood-oxygen-level-dependent signals were measured during optical self-stimulation of midbrain dopamine neurons in mice^24^; in contrast to the robust activations observed in dopamine terminal fields, no stimulation-related decreases were found. Moreover, the average baseline firing rate of dopamine neurons is less than 5 Hz^25^, and the maximal firing rate is thought to reach up to 50 Hz in vivo^26, 27^. If so, the number of inhibited neurons would have to greatly exceed the number that are directly stimulated in order to offset their activation. This makes the absence of reported inhibitions all the more striking.

### Relation to prior studies of intracranial self-stimulation

In an investigation of prediction-driven inhibition of dopaminergic neurons^27^ mice worked for stimulation of substantia nigra pars compacta dopamine neurons during the first portion of test sessions, and the sequence of stimulation trains recorded during the first phase was delivered non-contingently during a subsequent phase. The amplitude of stimulation-induced dopamine transients during the passive-playback phase was larger than during the initial phase, a result consistent with the prediction-driven inhibition of dopamine release postulated by the dopamine-RPE hypothesis. However, the stimulation-induced dopamine release did not decline preferentially over the course of the test sessions when the stimulation was self-administered.

Indeed, the decline was more reliable in the passive-playback condition, where there was no external event or consistent self-generated response to which a reward prediction could be anchored. This aspect of the results is inconsistent with the dopamine-RPE hypothesis, which predicts a progressive decline in dopamine signaling in response to repeated self-administered, but not passively received, unpredictably timed, rewards. Instead, this result suggests a cause unrelated to learning, such as change in excitation-release coupling.

Our interpretation of the present findings in terms of a rewarding effect of phasic dopamine firing is at odds with TDRL-based accounts of intracranial self-stimulation^2, 28^ but consistent with longstanding lines of work based on psychophysical inference^29, 30^, modeling of operant conditioning^31^, and information theory^32, 33^. On these views, animals performing conditioning tasks or foraging in the natural environment record a temporal map^34^ of events, their subjective impact (termed “reward intensity” in the self-stimulation literature), and the subjective costs^35, 36^ of reward-seeking actions. The incomes (rate of reward × reward intensity) from the available activities, net of costs, are computed from the temporal map, and behavior is partitioned between them accordingly^31^. Problematic assumptions about interval timing are avoided^37^.

A model developed within this tradition to account for electrical intracranial self-stimulation^30, 35, 38, 39^ has been applied with initial success to optical self-stimulation of midbrain dopamine neurons^30^. In this model, the impulse flow in the dopamine neurons induced by a pulse train (induced firing frequency × the number of optically activated neurons) is transformed into a record of reward intensity in the temporal map by a logistic function based on the one first measured in rats working for electrical stimulation of the medial forebrain bundle^40^. The induced dopamine firing acts as a primary reward, as it does in our interpretation of the present findings.

### The slow drift in stimulation effectiveness

In Experiment 2, time allocation to pursuit of the medium-strength stimulation grew systematically over the course of repeated testing in five of six rats; in three of these rats, an upward drift was also apparent in the case of the high stimulation strength. At least in part, this may have been due to continuing expression of Channelrhodopsin-2 and its progressive accumulation in the plasma membrane over the many weeks of testing. A higher density of Channelrhodopsin-2 would have increased the excitability of Ventral Tegmental Area (VTA) dopamine neurons to optical stimulation and added additional neurons to the stimulated population. The fact that time-allocation-versus-pulse duration curves shifted systematically leftward over the course of testing (Figure 4b) is consistent with such a mechanism.

Alternative mechanisms that boost the post-synaptic impact of the optically induced firing also bear consideration as do novel reinforcement-learning models. In one such model^41^, dopamine acts simultaneously as an RPE to slowly adjust an adaptable algorithm implemented in the prefrontal cortex and as a substitute for an external reward-related input that influences decisions on a rapid timescale. The role of dopamine as a reward on the short timescale seems consistent with our results, and an additional role in slow learning may have contributed to the slow drift in stimulation effectiveness. That said, we wonder how this model could accommodate both the slow drift and the rapid initial learning of the trial sequence. Detailed simulations and analysis will be required in order to assess how well this “meta-reinforcement-learning” model accounts for our data.

### An alternative role for dopamine

By treating response-contingent dopamine firing as a primary reward, we swim against the mainstream current: the view that such firing acts as an RPE. In contrast, our results and interpretation are consistent with a recent reformulation of the role of phasic dopamine firing in learning^42–44^. On that view, a burst of firing in midbrain dopamine neurons informs the animal that a significant event has occurred, triggering a retrospective search for the cause. The results of multiple new experiments carried out by the authors are interpreted to be consistent with their novel proposed function for phasic dopamine signaling and not with TDRL^42–44^. Moreover, they postulate that phasic dopamine signaling sums with the “the innate contribution to the meaning of a stimulus” in defining causal targets for learning. In this sense, natural rewards and phasic dopamine signaling are commensurable, a view consistent with the interpretation that optically induced dopamine transients acted as rewards in our experiments and not as RPEs.

## Conclusion

The causal role attributed to dopamine burst firing in the dopamine-RPE hypothesis predicts that reward-seeking behavior cannot remain stable in the face of repeated activation of dopamine neurons. In violation of this prediction, we report prolonged, stable performance in the face of repeated dopaminergic activation. These findings align well with the demonstration that activation of midbrain dopamine neurons failed to alter action policies in the two-step task^45^ and with the reinterpretation of phasic dopamine signaling in terms of retrospective causal attribution^42^.

## Methods

## 1 Subjects, housing, and ethical certification

Subjects were six TH-Cre, heterozygous, male, Long-Evans rats. In order to avoid excessive weight gain and possible renal and diabetic pathologies, they were fed a maintenance diet (Envigo #2014).

An inverse, 12 h light/12 h dark light cycle was maintained in the animal colony. Most of the animals’ sleep occurs while the lights are on, and this ensures that behavioral testing, done between the hours of 9 am and 8 pm, disrupts their sleep patterns as little as possible.

All methods were in accordance with guidelines established by Concordia University’s Animal Research Ethics Committee’s Terms of Reference (Certification of Ethical Acceptability #30000302) and the Canadian Council on Animal Care’s Guide to the Care and Use of Experimental Animals.

## 2 Surgical procedures and materials

### 2.1 Anesthesia and preparation

The animals underwent surgery when they had attained a body weight of approximately 400 g and were singly housed thereafter. Anesthesia was induced with an injection of a ketamine/xylazine mixture (87 mg/kg of ketamine mixed with 13 mg/kg of xylazine, administered IP). Once the animal was anesthetized, an SC injection of atropine sulfate (0.02 to 0.05 mg/kg) was given to reduce bronchial secretions.

“HypoTears” (Polyvinyl alcohol 1% (10 mg/1 ml)) was applied to the cornea for protection since atropine blocks lubricating secretions. After an anesthetic ointment (Xylocaine) was applied to the animal’s external auditory meatus, the animal was mounted in the stereotaxic frame using ear bars with blunt tips. The head was secured by hooking the incisors over the tooth bar. The animal’s snout was placed in a nose cone providing 800 ml O2/min and 2% isoflurane to maintain anesthesia.

The fur on the dorsal surface of the animal’s head was shaved and the underlying skin was disinfected with 70% isopropyl alcohol followed by chlorhexidine gluconate 4% (Hibitane). Lidocaine was applied to induce local anesthesia at the site of incision. The operation began after there was no observable reflex response to a toe pinch. An incision was made to expose the skull, and the edges of the wound were retracted by small clamps. Burr holes were drilled over the stimulation targets and pilot holes were drilled for the four to six jeweller’s screws that serve as anchors for the dental acrylic that secures the fiber optic probes. Using a stereomicroscope to provide visual guidance, an incision was made in the dura mater within each of the burr holes over the stimulation targets.

The animal rested on a heating pad (Sunbeam E12107-819) to maintain normal body temperature. A fluff underpad was placed between the ventral surface of the rat and the heating pad to prevent burns. The animal was given 5-10 ml/kg/hr of Lactated Ringer’s solution throughout the surgery.

### 2.2 Injection of viral vectors

Two small burr holes were drilled bilaterally over the VTA to provide access for a 26-gauge injector filled with the viral vector, RAAV5-EF1a-DIO-hChR2(H134R)-EYFP (UNC Vector Core). The injector tip was aimed at the following coordinates: AP −5.4 and −6.2; ML ± 0.7 (all coordinates are deviations from Bregma in mm). Two infusions of 1 µl were performed in each penetration at depths of 8.4 and 7.4 mm. Infusions were performed at a rate of 0.1 μl/min using a precision pump (Harvard Instruments), a 10 μl hamilton syringe (Hamilton Laboratory Products, Reno, NV) connected to the injector by surgical tubing. To allow for diffusion, the injector was left in place for 10 min following each infusion.

### 2.3 Implantation of fiber-optic probes

The tips of the optical implants were positioned at the following coordinates: AP = 5.8; ML = 0.7; DV = 7.7 for the left hemisphere and AP = 5.8; ML = 2.03; DV = 7.46 at a 10° angle for the right hemisphere. A representative histological section is shown in Figure S12.

The angled implant in one hemisphere provided space for implants to be placed bilaterally. The implant consists of a short section of step-index optical fiber (300 ± 6 µm core, 325 ± 10 µm cladding, NA = 0.39). One end of the fiber was terminated in a 2.5 mm steel ferrule with a roughened surface, held in place by dental acrylic applied at the end of surgery.

### 2.4 Pain management and post-surgery care

After surgery, the rats were placed in a shoebox cage mounted on a heating pad. To alleviate post-surgical pain, buprenorphine (0.05 mg/kg) was administered SC before the animals started to wake up. Penicillin procaine (0.3 ml) was injected SC after the surgery.

## 3 Behavioral procedures

### 3.1 Screening for optical self-stimulation

The subject was connected to an SLOC diode laser (Model BLM462TA-500FC) by means of a 300 μm core, 0.39 NA fiber-optic patch cable (Thorlabs M69L02, Newton, NJ, USA), an optical rotary joint (Doric Lenses, Model FRJ_1x1_FC-M3, Québec, QC, Canada), and a custom-built fiber-optic patch cord^46^ terminating in a steel ferrule that was mated to the ferrule on the rat’s head by means of a ceramic sleeve. Screening and testing took place in a 32 x 23 cm box with a metal grid floor and Plexiglas walls. The lever that triggered stimulation retracted when stimulation was given. Optical pulse trains were 1 s in duration and consisted of 5 ms (ELOP18, ELOP21, ELOP22, OP13) or 10 ms (OP18 and OP20) pulses. Optical power was set to 20 mW, and the pulse frequency was set to 40 Hz. The initial optical pulse trains were triggered by the experimenter. If the animal showed signs of approach (sniffing, forward locomotion, scanning movements of the head), attempts were made to shape these behaviors into lever pressing by providing progressively selective, contingent stimulation. Only subjects that could be trained to self-stimulate were retained for further testing.

### 3.2 Triadic-trial paradigm

The behavioral paradigm^38, 47, 48^ traces the trajectory of reward seeking over time, reveals whether behavior adjusts abruptly or gradually after changes in stimulation strength, and whether rats can learn to predict the sequence of trials (Figure 2). The latency to press the lever at the start of a trial reflects the payoff that the rat predicts; the higher the expected payoff, the shorter the latency. A difference across trial types in the initial latency to press the lever indicates the subject’s sense of how the states are related. The lever is extended for 240 s during each trial. To earn a stimulation train, rats had to hold down the lever for a cumulative work time of two s.

Depression of the lever accompanied by illumination of a light positioned over the lever. Each pulse train was 1 s in length and consisted of pulses five ms in duration. After each optical stimulation, the lever retracted for a “black-out” delay of 2 s wherein the trial clock stopped. Trials were arranged in repeating triads consisting of consecutive leading, test, and trailing trials. Stimulation strength (pulse frequency) was high on leading trials (56 Hz) and low on trailing trials (2 Hz) The pulse frequency for the test trials, held constant throughout the trial, was selected at random from a set consisting of 56 Hz, 2 Hz, and an intermediate value selected individually for each rat so as to support a mid-range allocation of time to lever depression. A session consisted of 15 triads (45 trials). Whereas the stimulation strength on the leading or trailing “bracket” trials was always the same and could therefore have been predicted by a trained rat, the stimulation strength on the test trial remained unpredictable until the rat had sampled the stimulation. Each trial was followed by a random inter-trial interval selected from a lagged exponential distribution with a minimum of four and a mean of eight s. The cage light flashed during the inter-trial-interval, indicating a trial boundary, but no information was provided that could indicate what the following trial type would be.

Each rat ran 30 to 32 sessions within the triadic-trial paradigm. The trial sequence was randomized afresh under program control (Matlab, R2022a, The Mathworks, Natick, MA) prior to each session. The temporal parameters of the optical stimulation were determined by a computer-controlled digital pulse generator. Experimental events and data acquisition were controlled by a custom-written computer programme (‘PREF3’, Steve Cabilio, Concordia University, Montreal, QC, Canada) and custom-built hardware (Dave Munro, Concordia University, Montreal, QC, Canada).

## 4 Data analysis

Lever-depression responses and delivery of optogenetic stimulation were analyzed in Matlab 2022a. The dependent measure was time allocation: the proportion of time that the rat was deemed to be working for the stimulation. Trial time was divided into periods when the lever was depressed (“holds”) and periods when it was released. Total work time included 1) the cumulative duration of hold times during a trial, and 2) release times less than 1 s. The latter correction was used because during very brief release intervals, the rat typically stands with its paw over or resting on the lever^47^. Therefore, we treat these brief pauses as work and subtract them from the total release time.

The state of the lever {up, down} was recorded every 0.1s, the period of the trial clock. Thus time-allocation can be measured over a series of trials by averaging the binary values recorded at every tick of the trial clock. Even over hundreds of trials, these measures are quite noisy. To smooth the data, we averaged the time-allocation measurements over 2s blocks, a time resolution that is sufficient to assess the stability of performance over the 240 s during which the lever was extended and armed during a trial.

A set of piecewise linear functions was fitted to the time-allocation time courses. The set of functions consisted of 1) a single straight line, 2) a linear segment followed by a horizontal segment, 3) a horizontal segment followed by a linear segment, and 4) a horizontal segment followed by a linear segment followed by a second horizontal segment. The mkpp function in Matlab (R2022a, The Mathworks, Natick, MA) was used to create the piecewise linear functions, the ppval function to evaluate them, and the fit function to determine the best-fitting parameter values. The four functions differ in the number of parameters. Thus, the Akaike Information Criterion^49^, which penalizes candidates on the basis of their parameter number, was then used to select the best-fitting function.

The stability of time allocation across test sessions was assessed by fitting a linear function to the average within-session time-allocation for each test-trial stimulation strength as a function of session number and also of session date. (Five trials were run at each test-trial stimulation strength in each session. Thus these averages were computed across all timepoints and all five trials for each stimulation strength.) The time-course analysis revealed that time allocation varied systematically over the course of the trials (Figs 2e,f, S2,S3). Thus some of the variance around the average within-session time-allocation for a given trial type is due to the systematic evolution of time allocation as a function of trial time. To isolate the noise variance, we used the best-fitting time-course function to back out the contribution of the within-trial time course. The resulting residual variance was used to compute the bootstrapped 95% confidence intervals shown in Figures 4a and S9,10. These figures show a systematic upwards drift in time allocation over the course of repeated testing. Thus, some of the across-session variance in time-allocation at any time point was due to this drift. To isolate the noise variance, we used the best fitting linear function fitted to the within-session averages for each test-trial stimulation strength to back out the effect of the drift. The resulting residual variance was used to compute the bootstrapped 95% confidence intervals shown in Figures 2e,f and S2,3.

The latency of the first lever depression on each trial and the inter-press interval were extracted and pooled across sessions by trial type. The bootstrapped 95% confidence intervals around the latencies were also calculated and plotted as cumulative distributions.

All analyses were done individually for each of the rats in both experiments as well as for each set of simulated data.

## 5 Simulations

The simulations are accessible at the following URL: https://github.com/carterfrancis/RandomWorld_Simulations.

The agent used State-Action-Reward-State-Action (SARSA), an on-policy algorithm implementing one-step Temporal-Difference learning, TD(0). This learns q values, estimates of true action values, through linear function approximation.

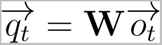

where q(t) is the vector of estimated action values at time t, W is the matrix of weights and o(t) is the one-hot state vector. q(t) contains action values for all possible action-state pairs, including work and leisure activities (resting, exploratory sniffing, and grooming). An action a_i_(t) is selected from q(t) using a combination of softmax (with temperature parameter of 1) and - greedy (є = 0.9) action selection to allow for gradual changes in the level of pressing and to prevent saturation/maximal pressing. This is because the behavior of the rats saturated at time allocation values below 1. Thus, the probability of selecting action a_i_ is

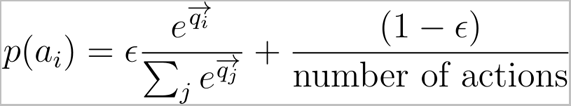

At each moment, the agent selects an action according to these probabilities and the plotted time allocation comes from the proportion of trial times when the agent selected the pressing action (error bars are the 95% CI of the mean).

Weight matrix updates used the RPE, *δ*, to indicate the correct amount of change. More specifically, the agent used differential SARSA to make use of the average-reward definition of delta, the RPE. The average-reward definition of RPE was used because it is appropriate for continuing tasks and avoids problems associated with discounting.

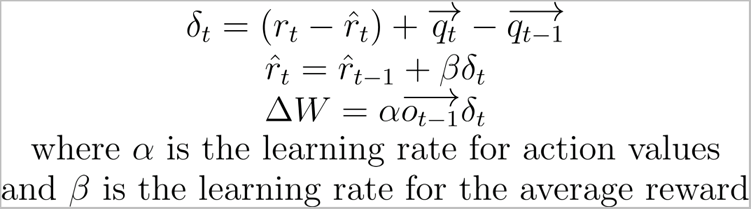

### 5.1 Modeling the paradigm

Simulations for figure 2 were carried out for individual trials run at different stimulation strengths. In the “DA=RPE” simulation, the pressing action set delta to the strength of stimulation. In the “DA=reward” simulation, the pressing action provided a reward (r_t_), and the equations above determined the RPE.

In figure 3, we simulated the entire behavioral paradigm, including alternating trial types, with stimulation eliciting a reward (DA=Reward). We altered the correspondence between true states and the states provided to the agent, to reflect different degrees of understanding of the paradigm. In the *complete state information* simulation, the rat’s state vector necessarily followed the triadic structure of trials. To designate this perfect parsing out of situations, we can give the rats 6 possible states: 1-the rat is in test trial and hasn’t sample the lever yet 2-3-4-rat is in test trial with knowledge of low,med,hi stimulation strength, respectively 5-the rat is in a leading trial, 6-the rat is in a trailing trial. In the *incomplete state information* simulation, the rat does not have a memory that can look back more than one trial, and comes to expect an alternation of high and low trials. The same vector is used as in the *complete information* simulation, but on half of transitions between trials, the rat believes it is entering a high trial following a low, or a low following a trial worth pressing for (high or med).

Figure 2c (DA=Reward) corresponds to the test trial pressing in the *complete state information* simulation. Figure 2d (DA=RPE) corresponds to individual and independent trials where the rat learned action values for that single trial (one state). This was done because simulating the entire paradigm with DA=RPE led to saturation of the action values much too early, so the average over many triads corresponds to maximal pressing and does not show the growth which we see in individual trials.

## 6 Fast-scan cyclic voltammetry

### 6.1 Subject

The subject was a male Long-Evans TH-Cre^+/-^ rat maintained on a 12-hour reverse light/dark cycle (lights off from 08:00 to 20:00) with ad libitum access to food and water.

### 6.2 First surgery: virus injection and implantation of optical implant

The rat was anesthetized with a mixture of ketamine hydrochloride (87 mg/kg) and xylazine hydrochloride (13 mg/kg) (injected i.p.). It received a s.c. injection of atropine sulphate (0.05 mg/kg) prior to surgery to reduce bronchial secretions, an s.c. injection of buprenorphine to alleviate post-operative pain (0.05 mg/kg), and an s.c. injection of penicillin procaine to prevent infection (0.3 cc). Burr holes were made to provide access for the injector above the VTA.

VTA DA neurons in TH::Cre males were infected with a serotype-5 adeno-associated virus (AAV5) containing an EF1a-DIO-hChR2(H134R)-EYFP-WPRE transcript suspended in phosphate-buffered saline. The rat received six injections of 0.5 µL in each hemisphere (AP: 5.4, ML: 1.0, DV: 8.3, 7.7, 7.2; AP: 6.2, ML: 1.0, DV: 8.3, 7.7, 7.2), delivered at a rate of 0.1 µL per min via a Hamilton syringe connected to a Harvard infusion pump. The injections were separated by a 10-min waiting period.

An optical implant was lowered into each hemisphere at a 10-degree angle along the medial-lateral axis and aimed at the following coordinates: AP 5.8, ML 2.2 and DV 8.12. The optical implant was fashioned from an optical fiber (300 µ core diameter, multimode, step-index, 0.39 NA, Thorlabs FT300UMT), stripped of its cladding. One end was inserted in a stainless alloy ferrule, glued in place and polished. The optical implants were secured with dental acrylic and anchored with jeweler screws.

The rat rested on a heating pad throughout the surgery. An s.c. injection of Ringer’s solution was given every hour to maintain hydration.

### 6.3 Second surgery: implantation of chronic carbon-fiber microsensor and reference electrode

Burr holes were made bilaterally above the nucleus accumbens, and the carbon-fiber microsensors were lowered, one at a time. The microsensors were fashioned as described by Clark, SandBerg, Wanat et al. (2010). One carbon fiber was pushed through a section of silica tubing while immersed in alcohol and was subsequently allowed to dry. Then, the microsensor was sealed with resin epoxy and fixed to a gold-plated Amphenol connector with silver epoxy. The sensors were pre-cleaned with a solution of 2-propanol containing activated carbon. The were aimed at the NAc shell (AP: 1.7 mm; ML: 1.0 mm; DV: 7 mm from the skull) and lowered slowly, by 0.2 mm every 30 s, to minimize damage.

An Ag/AgCl reference electrode was fashioned from a silver wire secured with silver epoxy to a gold-plated Amphenol connector and soaked in a 10% solution of sodium hypochlorite for 60 mins. The final conductive surface was approximately 5 mm in length. The reference electrode was placed in the left hemisphere posterior to the head cap built during the first surgery. The location of the tip was roughly AP: −9 mm; ML: 4 mm; DV 5 mm from bregma.

### 6.4 Behavioral testing

Behavioral procedures were as described above (Section 3), except that the black-out delay was set to 8 s to ensure that the full voltammetric response to the stimulation train could be acquired. Optical pulse trains were generated by a computer-controlled Master-8 pulse generator (A.M.P.I., Jerusalem, Israel) connected to a 473 nm DPSS laser (Laserglow, Toronto, Ontario). The output of the laser was routed to the chronically implanted optical probe via an optical rotary joint (Doric Lenses FRJ_1x1, Quebec, Quebec) and a custom-built optical patch cord^46^. A custom-written computer program (“PREF”, Steve Cabilio, Concordia University, Montreal, QC, Canada) controlled the experiments and logged the behavioral data.

### 6.5 Electrochemistry

Cyclic voltammograms were generated at 10 Hz by applying an 8.5 ms triangular waveform that ramped from -0.4 V to +1.3V and back to -0.4 V at a scan rate of 400 V/s. The potential was held at -0.4 V between scans to promote cation absorption at the surface of the FSCV electrode. The triangular waveform was generated using an analog output of a PCI-6052E multifunction data-acquisition / digital-analog converter board (National Instruments, Austin, TX) controlled by LabView (National Instruments, Austin, TX). Generation of the triangular waveform was gated by a PCI-6711E board (National Instruments, Austin, TX).

All potentials were measured with respect to the Ag/AgCl reference electrode using a headstage based on a design by Scott Ng-Evans (University of Washington). The output of the headstage was connected to an analog input of the PCI-6052E board to monitor the current passing through the FSCV electrode.

The FSCV data were logged by software written by M. L. A. V. Heien and R. M. Wightman (Department of Chemistry, University of North Carolina, Chapel Hill, NC, USA). The PREF system outputted a signal to a digital input of the PCI-6711E board that allowed the FSCV software to timestamp and record the onset of each stimulation train.

### 6.6 Analysis of electrochemical data

Faraday currents registered by chronically implanted microsensors evolve over time as the FSCV electrode interacts with the brain environment and is etched by continuous usage (Keithley, Takmakov, Bucher et al., 2011; Rodeberg, Johnson, Cameron et al., 2015). Consequently, calibration data were derived from measurements acquired in awake animals presented with unexpected stimulation trains.

The Principal Component Regression (PCR)-Principal Component Analysis (PCA) methods used in many voltammetry experiments^50^ were employed to estimate peak-dopamine concentrations and pH changes. Background subtraction was omitted to provide increased predictive power^51^ while avoiding the arbitrariness entailed in determining what constitutes the background and distortions that can arise from attempts to remove it. A relatively large proportion of principal components was retained so as to include all informative parts of the voltammograms^51^. Although the full method developed by Kishida et al.^51^ would have been even more powerful, the number of voltammograms required was unfeasible, and we settled on a compromise that economized on the number of unexpected stimulation trains delivered to acquire the training set.

The training sets were acquired from six rats presented with unexpected, experimenter-triggered optical pulse trains delivered at different pulse frequencies (optical power: 40 mW; pulse frequencies: 5, 10, 20, 40, and 56 Hz). The selection of cyclic voltammograms was based on established features of the cyclic voltammograms obtained from microsensors exposed to rapid changes in DA or pH^52^. Due to the high degree of similarity across microsensors, data from all six animals were combined in the same training set. A single matrix of cyclic voltammograms was thus constructed (Matrix **A**: n x 1000; number of samples by number of voltages tested for each scan) as well as a second containing the corresponding changes in DA concentration and pH (Matrix **C**: n x 2; number of analytes {[DA], pH} by number of samples).

To build a linear model to predict DA concentrations and pH changes from cyclic voltammograms obtained from solutions containing known DA concentrations at known pH, we solved for:

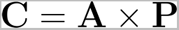

where **A** is the matrix containing the voltammograms, **C** is the matrix containing the known concentration associated with these voltammograms for both analytes, and **P** is the regression-coefficient matrix.

Before conducting the regression, we used PCA to reduce the dimensionality of matrix A. This step lowers the number of samples of known concentration required to obtain a statistically reliable model. Because each voltammogram consists of 1000 data points, solving **C** = **PA** would require several thousand scans with known concentrations. Such numbers can be reached with in vitro calibration data but obtaining that large a number in awake animals is unfeasible.

Principal components were calculated using singular-value decomposition in MATLAB. A scree plot was used to determine the number of components that would account for at least 90% of the variance in matrix A. Fifteen components were retained. We expressed the voltammograms as a function of the 15 dimensions determined by PCA to obtain matrix **A_pro_**_j_.

We solved for P using the least squares method:

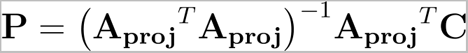

where **A_proj_^T^** is the transposition of matrix **A_proj_**

The coefficient matrix, **P,** could then be used to predict DA concentrations and pH changes from the voltammograms acquired during triadic-trial testing.

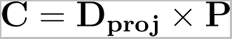

where **D_proj_** is the data expressed as a function of the 15 dimensions determined by PCA and **C** is a matrix predicting DA concentration and pH during the trials.

The number of components retained greatly exceeds the two or three recommended when data are background subtracted^50, 53^. However, retaining such a large number of components is consistent with the observation of Kishida et al.^51^ that voltages obtained at multiple time points during the scan have predictive power to detect and quantify DA transients.

## Acknowledgements

This work was supported by a grant from the Natural Sciences and Engineering Research Council of Canada to Peter Shizgal (RGPIN-2016–06703). Marie-Pierre Cossette and Vasilios Pallikaras were supported by Alexander Graham Bell Canada Graduate Scholarships (CGS D). Ivan Trujillo-Pisantry was supported by a MELS-PBEE scholarship from the Fonds de la recherche en santé de Québec (#140478) and a scholarship from Consejo Nacional de Ciencia y Tecnologia (CONACYT, #309126, Mexico).

Peter Dayan provided invaluable advice, constructive critique, and encouragement at all stages of this project. He has been exceptionally generous in his support and open to considering views that differ from his own.

The authors thank Matt Bovinick, Zeb Kurth-Nelson, and Jane Wang for their very helpful consultation about the applicability of their meta-reinforcement-learning model^41^ to our data.

David Natanael Velázquez Martínez contributed to troubleshooting the methods for Experiment 2 and to training several of the subjects.

The FSCV data presented here as well as the method for estimating dopamine concentrations appeared originally in Marie-Pierre Cossette’s doctoral thesis^54^.

Stephen Cabilio developed, adapted, and maintained the experimental-control and data-acquisition software for the behavioral experiments.

The experimental-control and data-acquisition hardware was designed, built, and maintained by David Munro.

Christopher Law provided helpful advice and assistance with microscopy and image processing. Éric Borduas helped prepare the video.

Karl Deisseroth and Ilana Witten donated TH::Cre sires to establish our breeding colony.

## Data-accessibility statement

The raw data have been uploaded to the public-access repository at the Open Science Foundation: https://osf.io/uw3nb/.

## Abbreviations

DA: Dopamine
ChR2: Channelrhodopsin-2
fMRI: functional Magnetic Resonance Imaging
FSCV: Fast-Scan Cyclic Voltammetry
PCA: Principle Component Analysis
RL: Reinforcement Learning
RPE: Reward Prediction Error
TD: Temporal Difference
TDRL: Temporal Difference Reinforcement Learning
VTA: Ventral Tegmental Area

## Supporting Information

**Figure S1:**
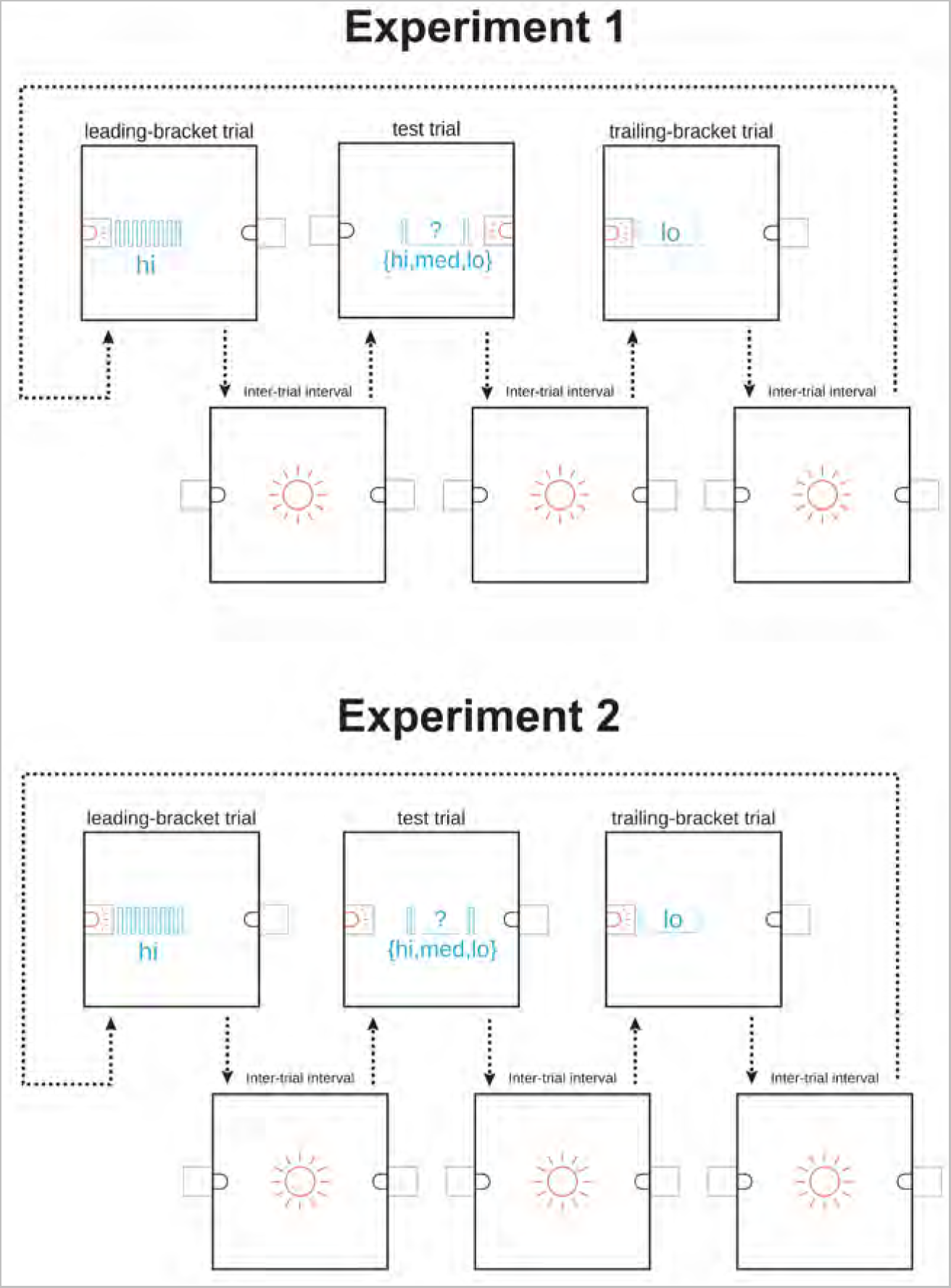
Testing paradigms. In Experiment 1, different levers were used to trigger stimulation delivery on bracket (upper left and right panels) and test (upper middle panel) trials. Stimulation strengths (pulse frequencies) are shown in cyan. The paradigm employed in Experiment 2 is identical to the one employed in Experiment 1 with one exception: the same lever was used to trigger stimulation delivery on bracket (upper left and right panels) and test (upper middle panel) trials.

**Figure S2:**
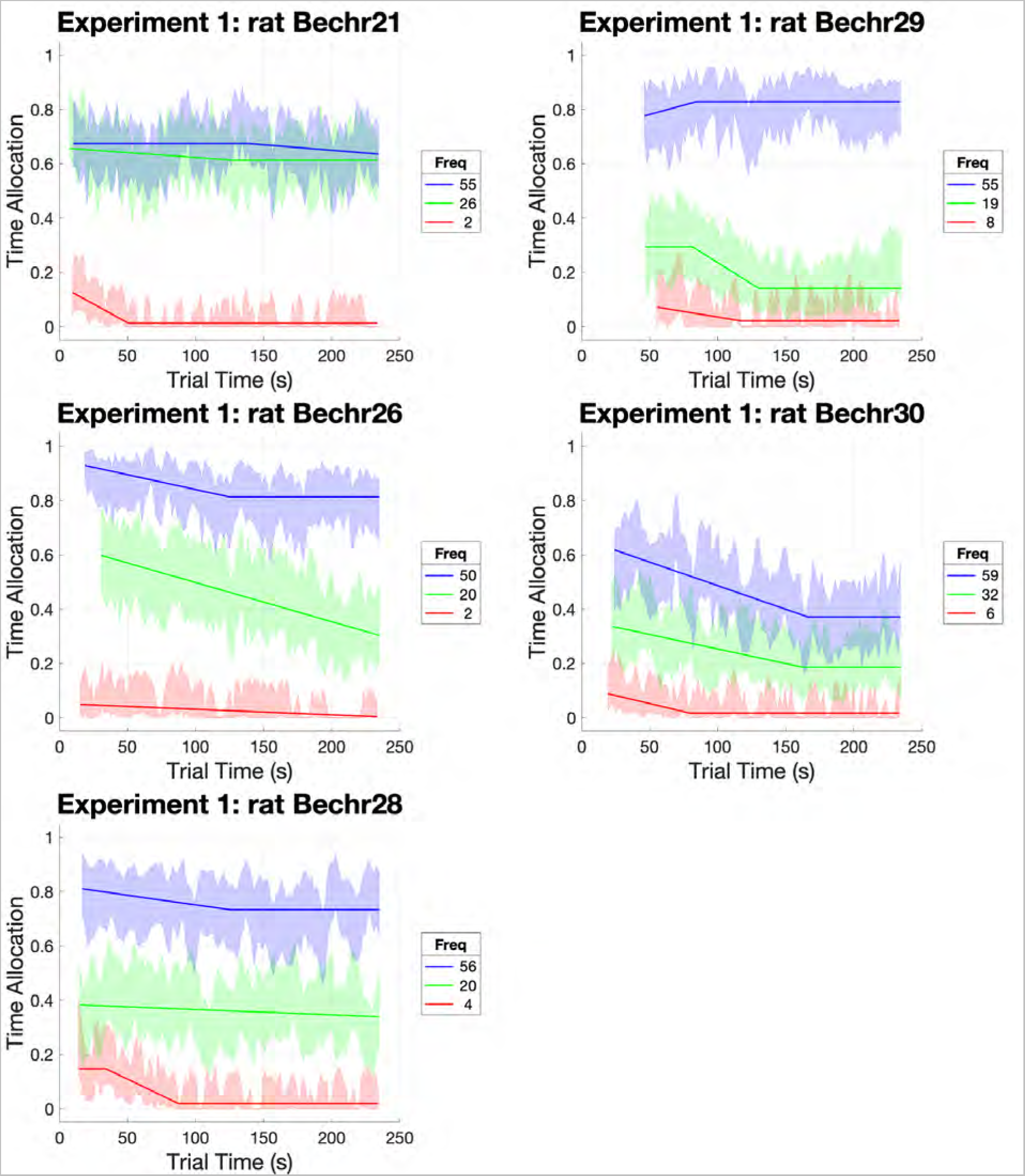
Stability of test-trial responding for rewarding optical stimulation for each subject in Experiment 1. The proportion of time that the lever was depressed is shown in successive 2-s time blocks. Lines show best-fitting piecewise linear functions, and error bands show the surrounding 95% confidence intervals. The stimulation strength was set by the pulse frequency, which was either high, medium, or low. The plots for each stimulation strength begin at the time block when responding had commenced on 75% of the trials and end at the 3rd to last time block.

**Figure S3:**
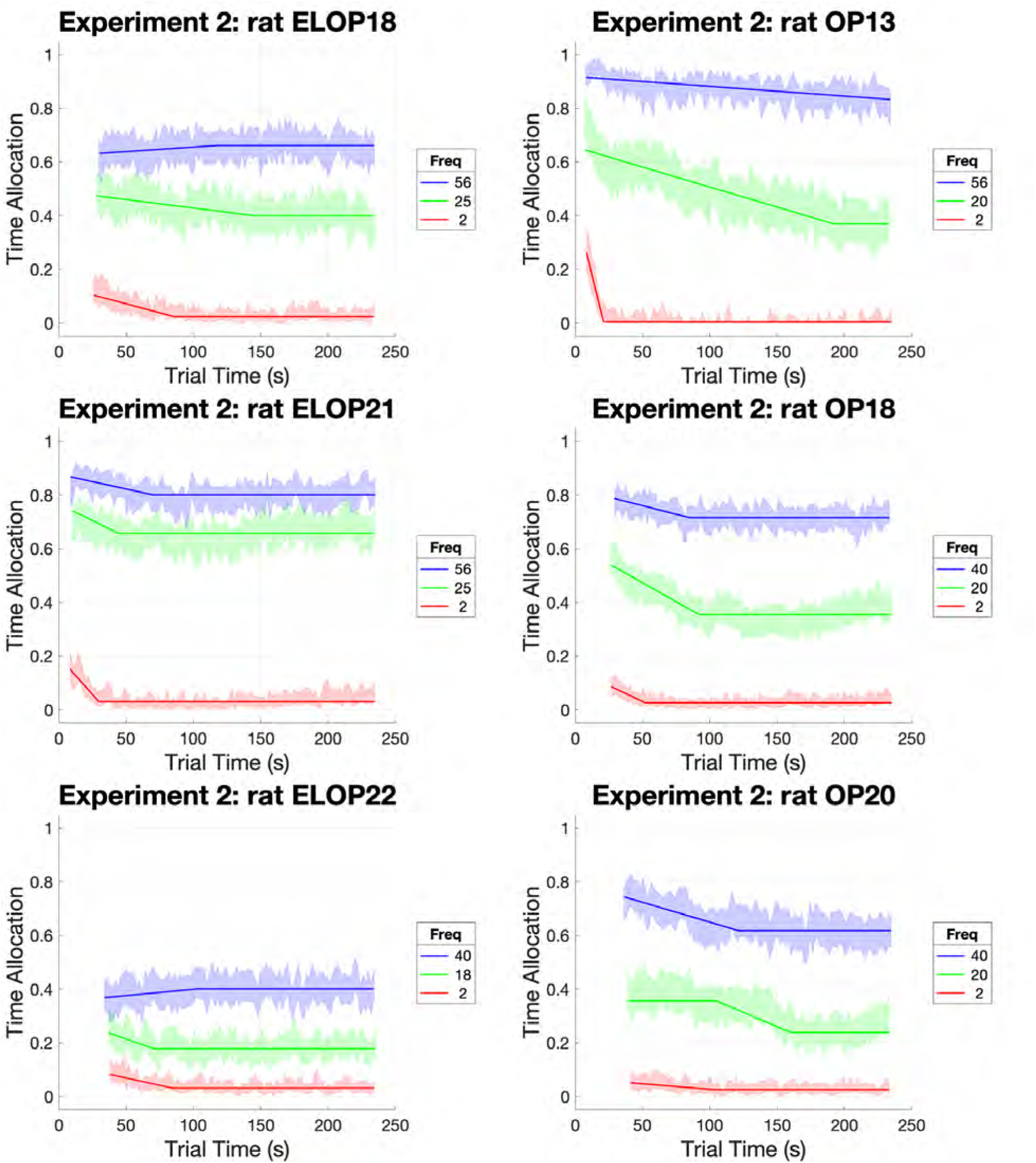
Stability of test-trial responding for rewarding optical stimulation for each subject in Experiment 2. For details, see the caption for Figure S2.

**Figure S4:**
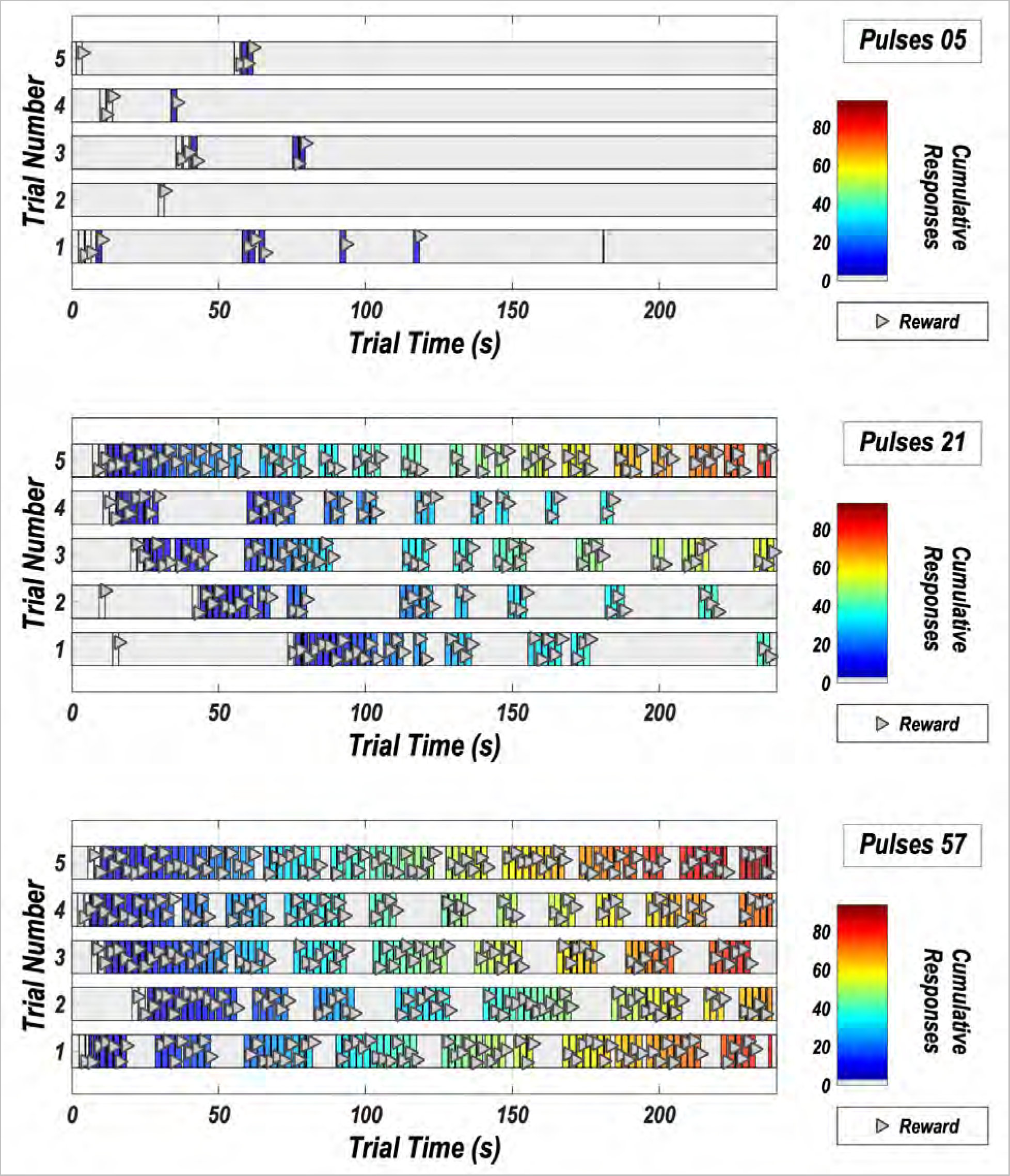
Pattern of responding in Experiment 1. The data are from rat Bechr28, test session 3. Responding for the low, medium and high stimulation strength is shown in the upper, middle, and bottom panels, respectively. Bout durations, and response totals (color map) typically increase as a function of increasing stimulation strength, whereas inter-bout pauses typically decrease. The triangles indicate lever presses and are jittered vertically to increase visibility.

**Figure S5:**
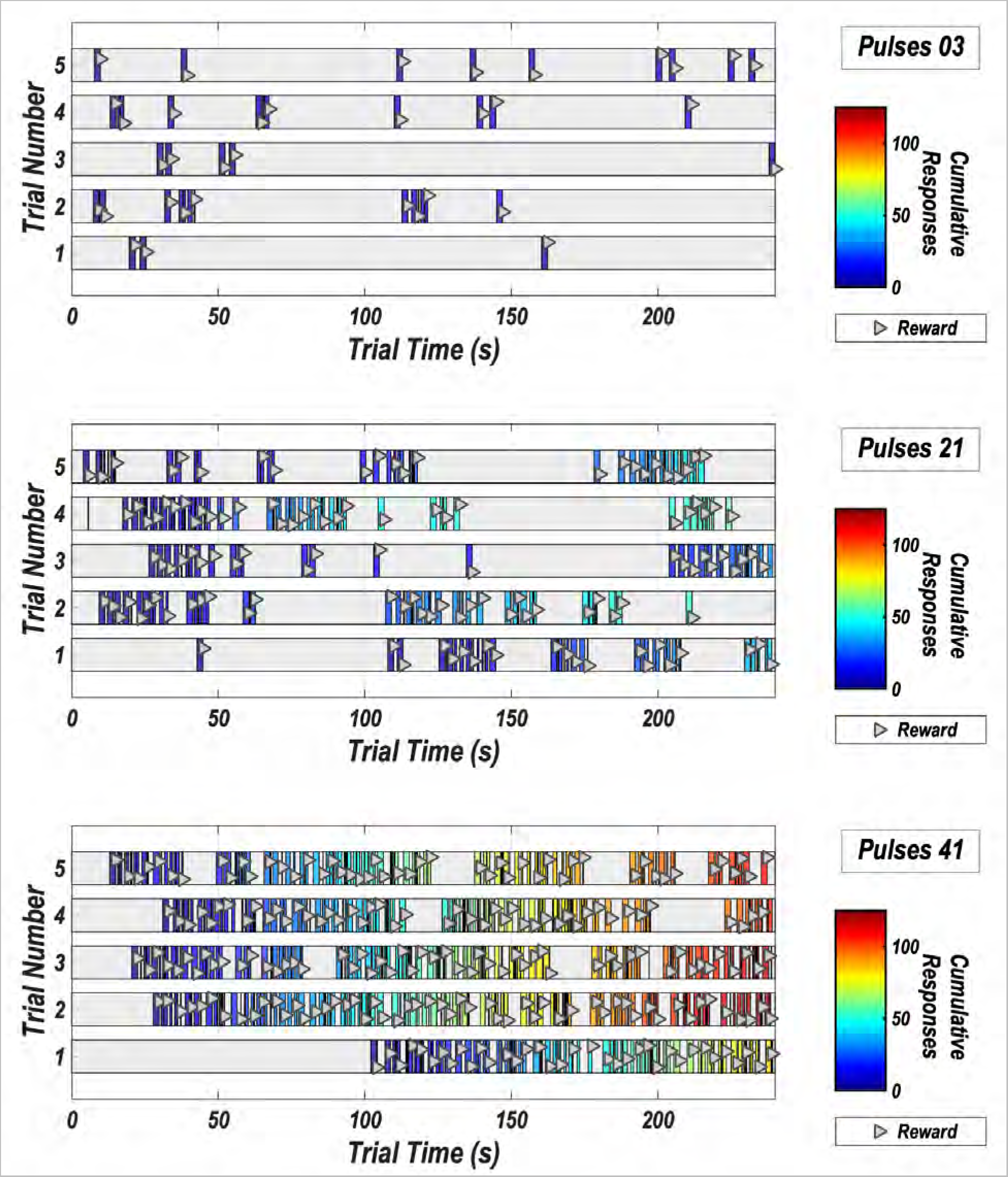
Pattern of responding in Experiment 2. The data are from rat OP18, test session 5. Responding for the low, medium and high stimulation strength is shown in the upper, middle, and bottom panels, respectively. Bout durations, and response totals (color map) typically increase as a function of increasing stimulation strength, whereas inter-bout pauses typically decrease. The triangles indicate lever presses and are jittered vertically to increase visibility.

**Figure S6:**
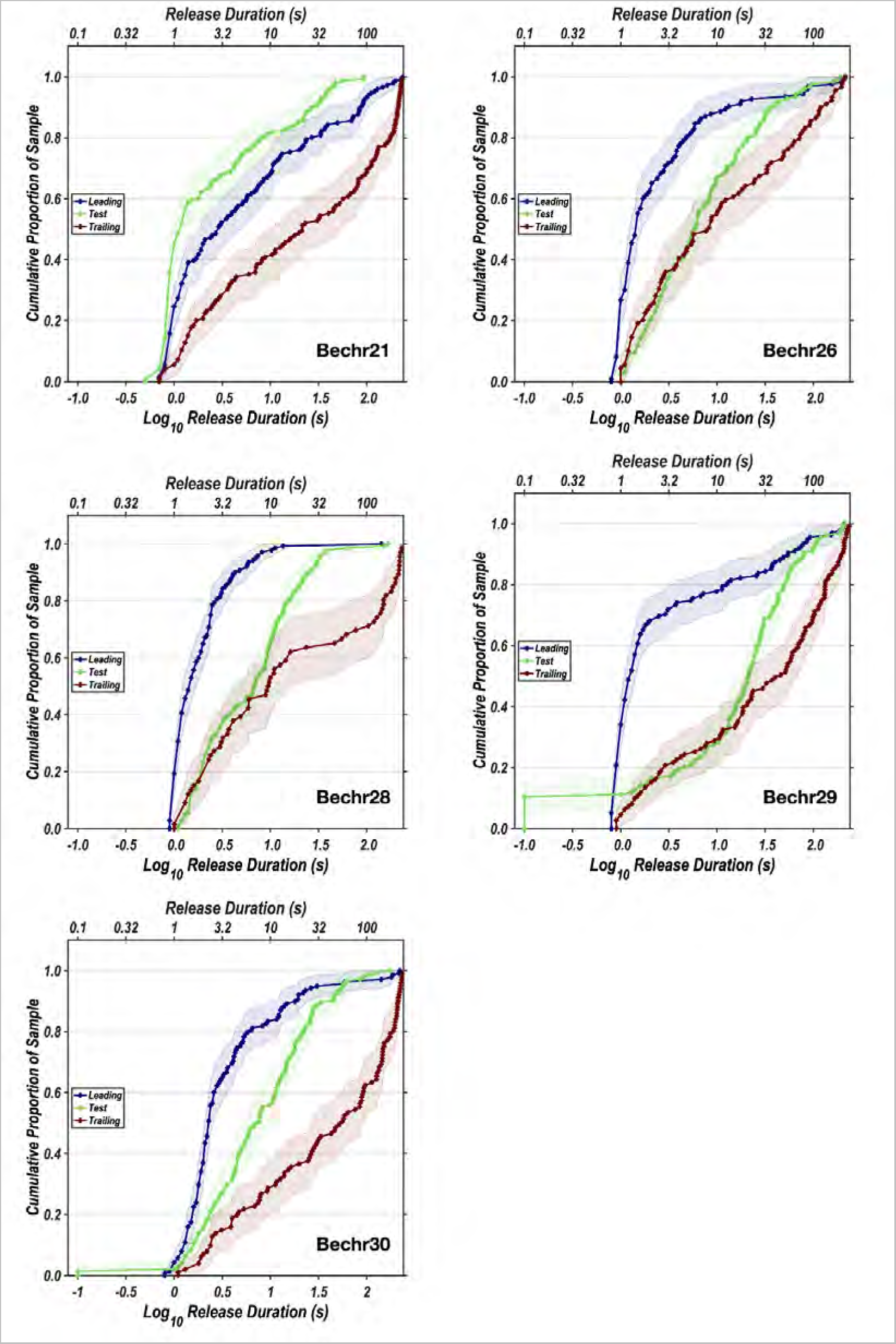
Cumulative distributions of the initial latencies on all trial types in Experiment 1.

**Figure S7:**
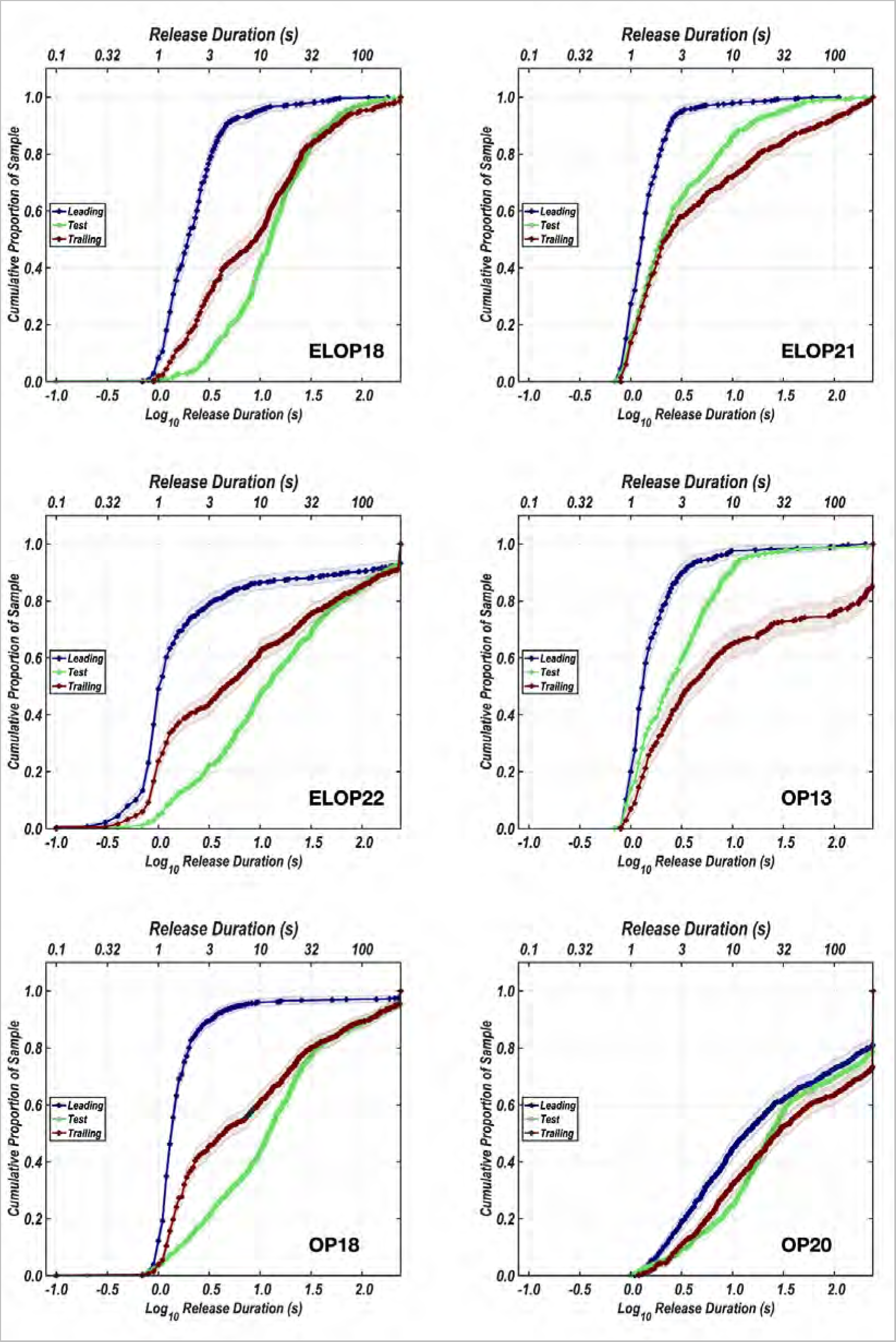
Cumulative distributions of the initial latencies on all trial types in Experiment 2.

**Figure S8:**
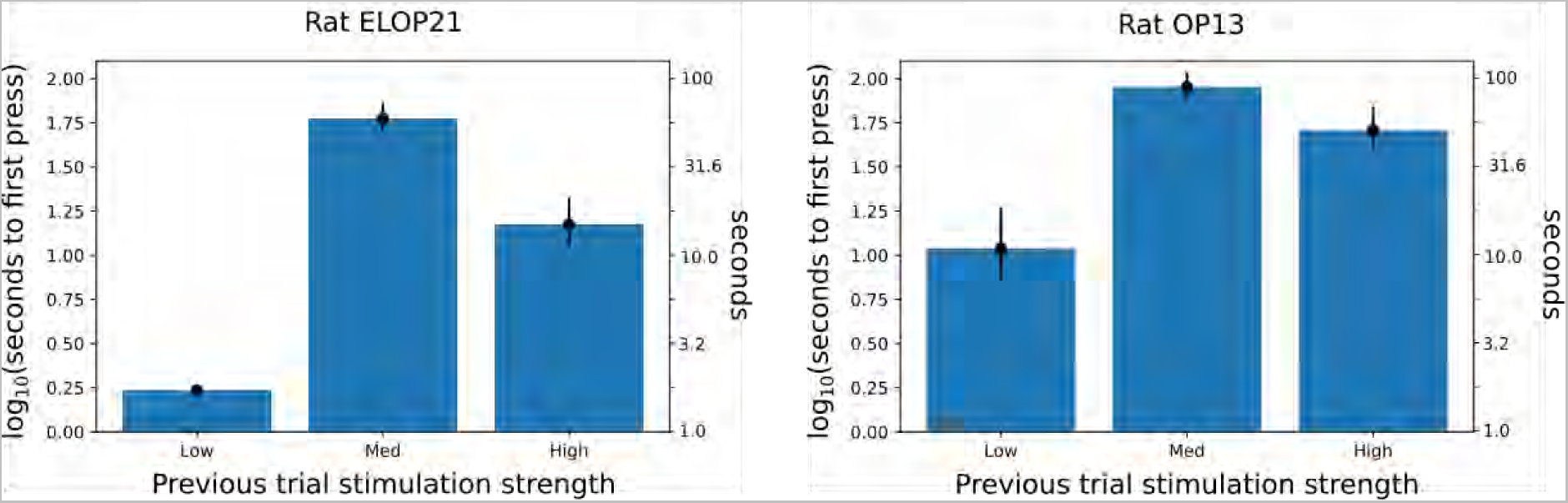
Bar plot of latencies until first press on trailing trials depending on the stimulation strength on the previous test trial. Data are from the two rats that learned the entire triadic trial structure.

**Figure S9:**
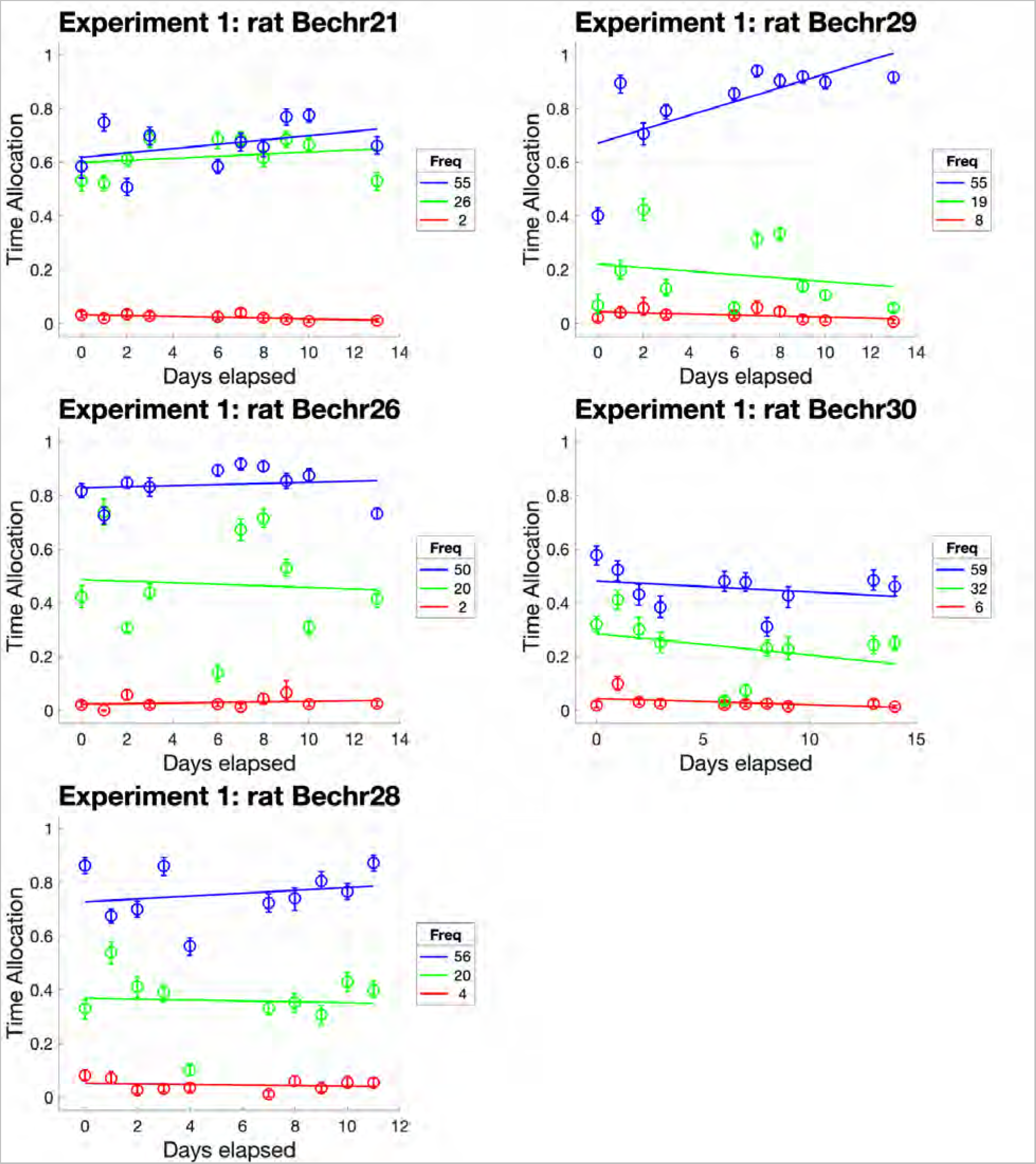
Evolution of time spent working as a function of stimulation strength over the course of Experiment 1. Many more sessions were run in Experiment 2, thus extending the period over which drift could be observed.

**Figure S10:**
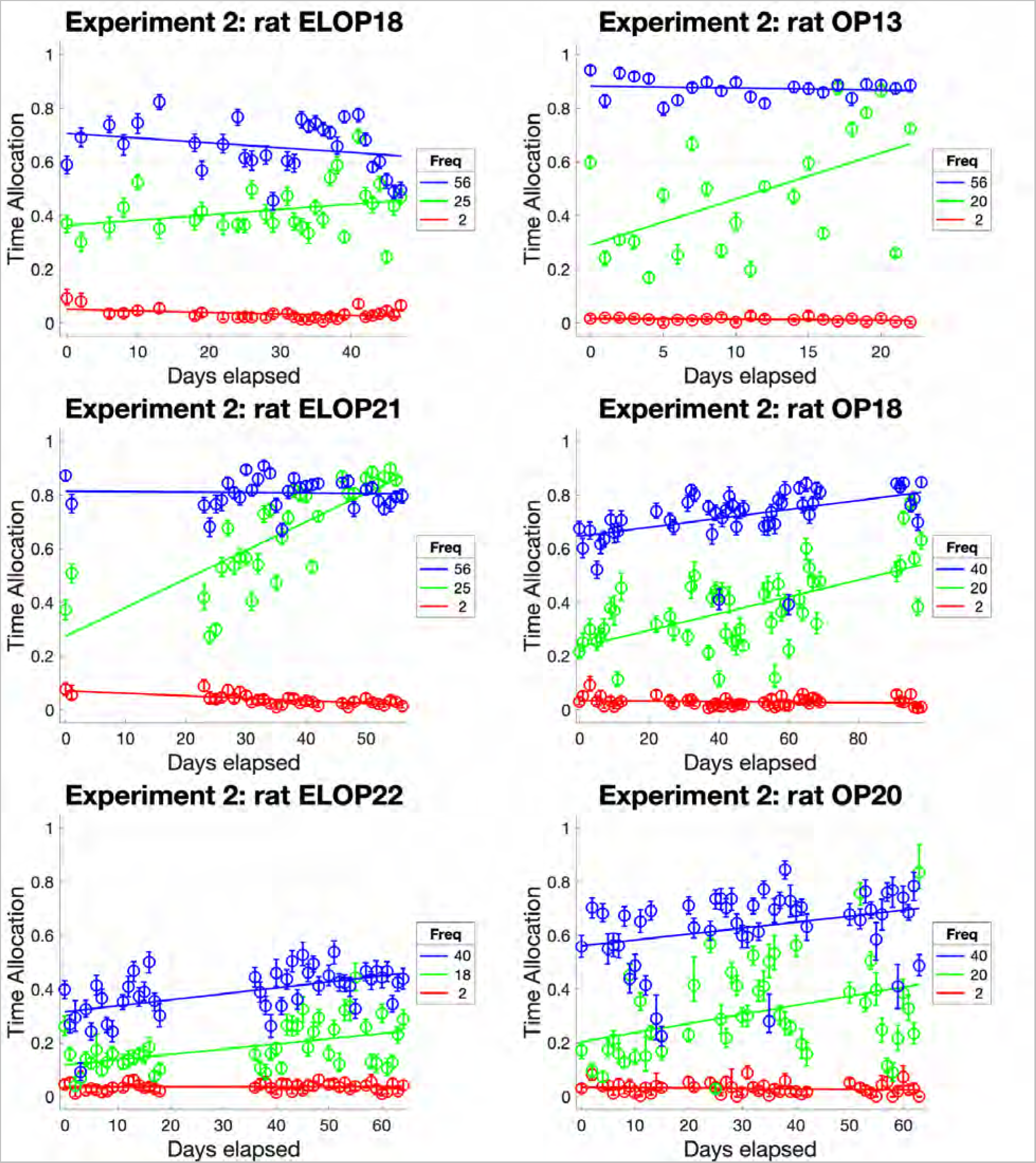
Evolution of time spent working as a function of stimulation strength over the course of Experiment 2. Over the prolonged period of repeated testing, time allocated to working for the medium-strength reward increased reliably in all rats, save for ELOP18. Data points are means over the five trials carried out at each stimulation strength during each test session.

**Figure S11.**
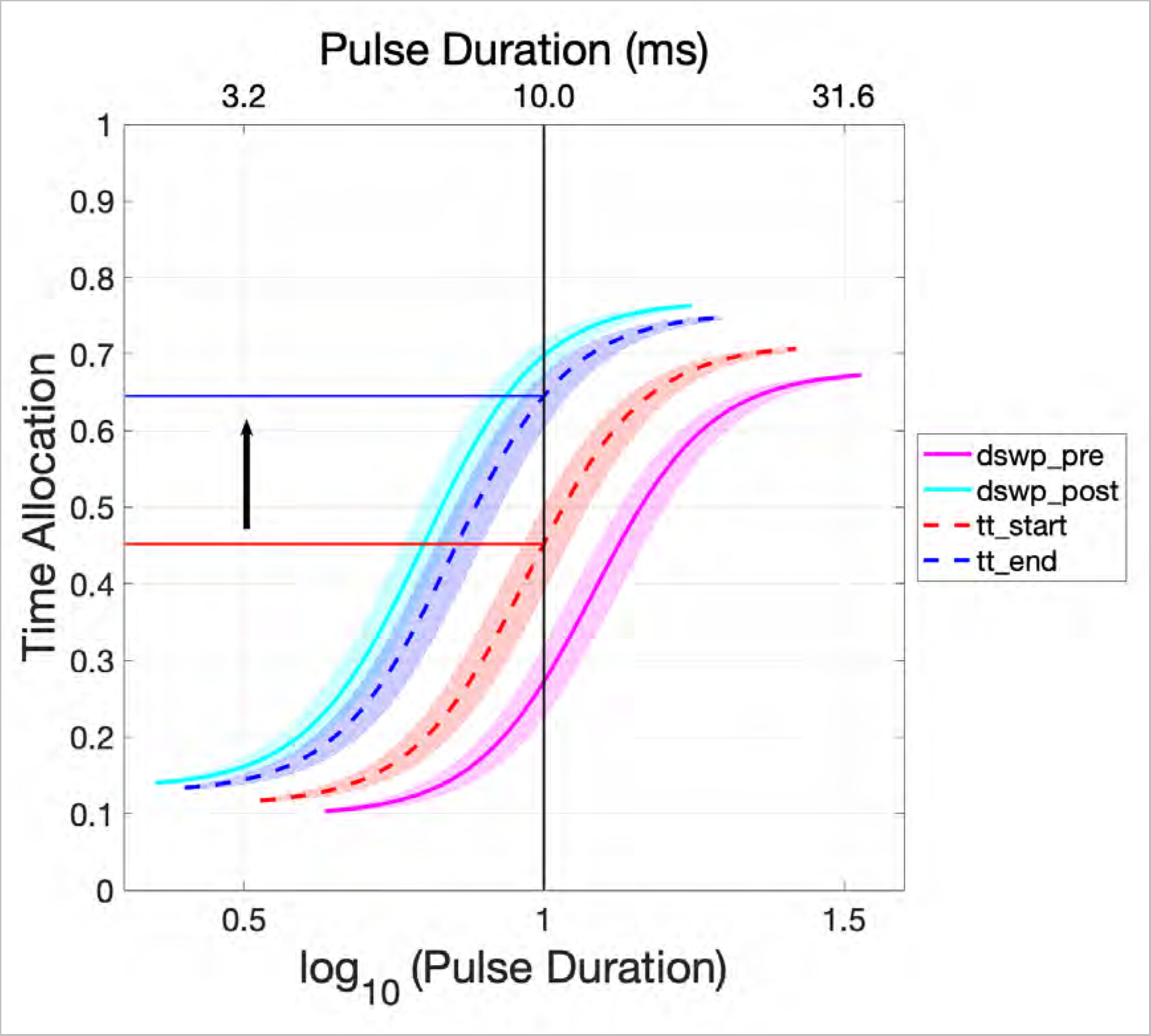
Fitted and interpolated sigmoidal functions. The solid curves represent sigmoidal functions fitted to pulse-duration sweeps obtained from rat OP18 prior (dswp-pre) and after (dswp-post) triadic-trial testing. The dashed curves are interpolated sigmoids for the start (tt-start) and end (tt-end) of triadic-trial testing. These were generated on the basis of the slope of drift in Figure 4b, the start- and end-dates of triadic-trial testing, and the parameters of the sigmoids fitted to the data from the pre- and post-triadic-trial pulse-duration sweeps. Shaded areas denote the 95% confidence intervals surrounding the position-parameter estimates. The solid blue and red lines show the predicted change in time allocation over the course of triadic trial testing, which was carried out at a pulse duration of 10 ms.

**Figure S12.**
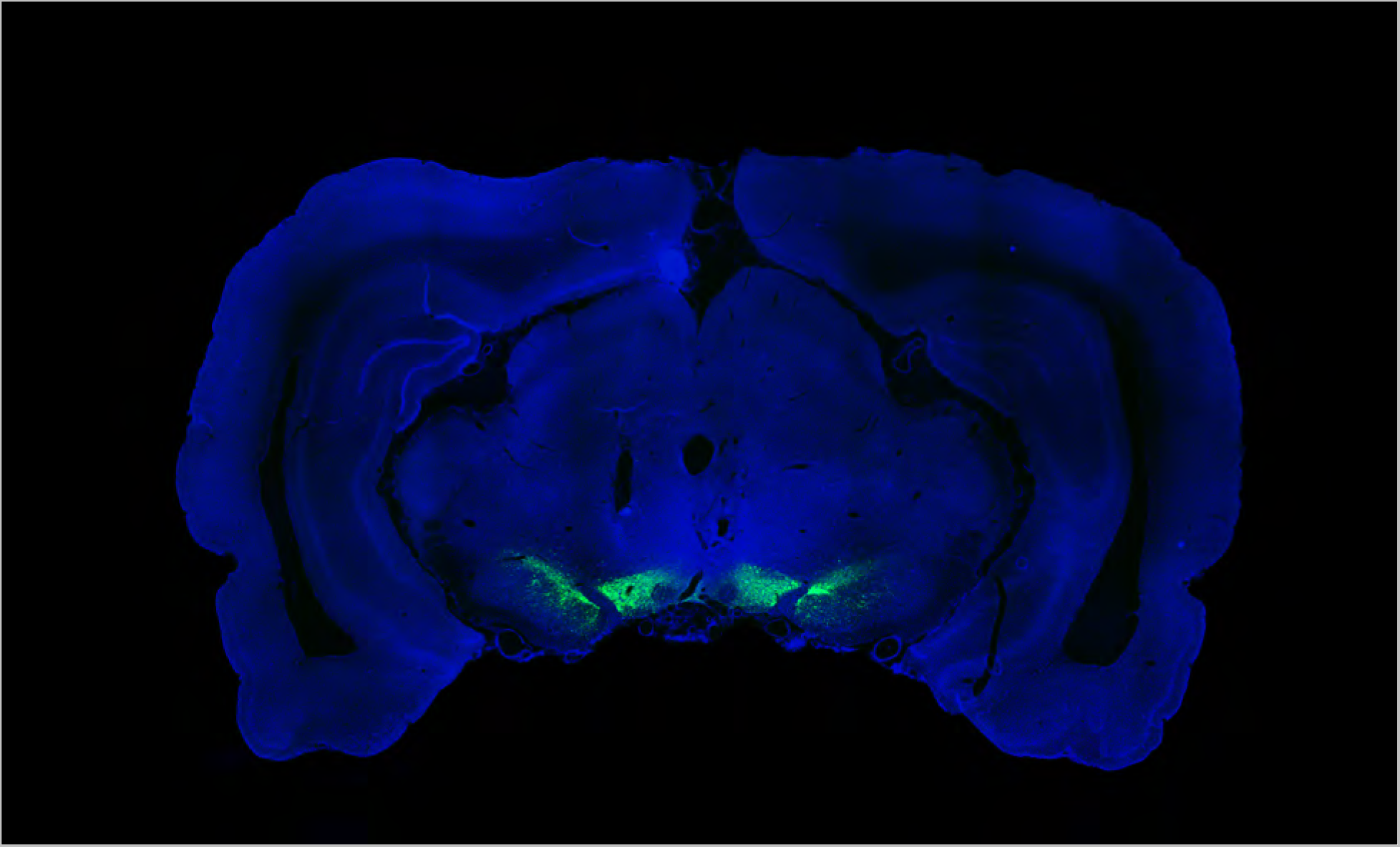
Histological image for subject OP13 (Experiment 2); eYFP expression (green) shown along with DAPI (blue) for anatomical reference (image enhanced by rescaling the distribution of pixel intensities). This figure was published previously^1^.

**Figure S13.**
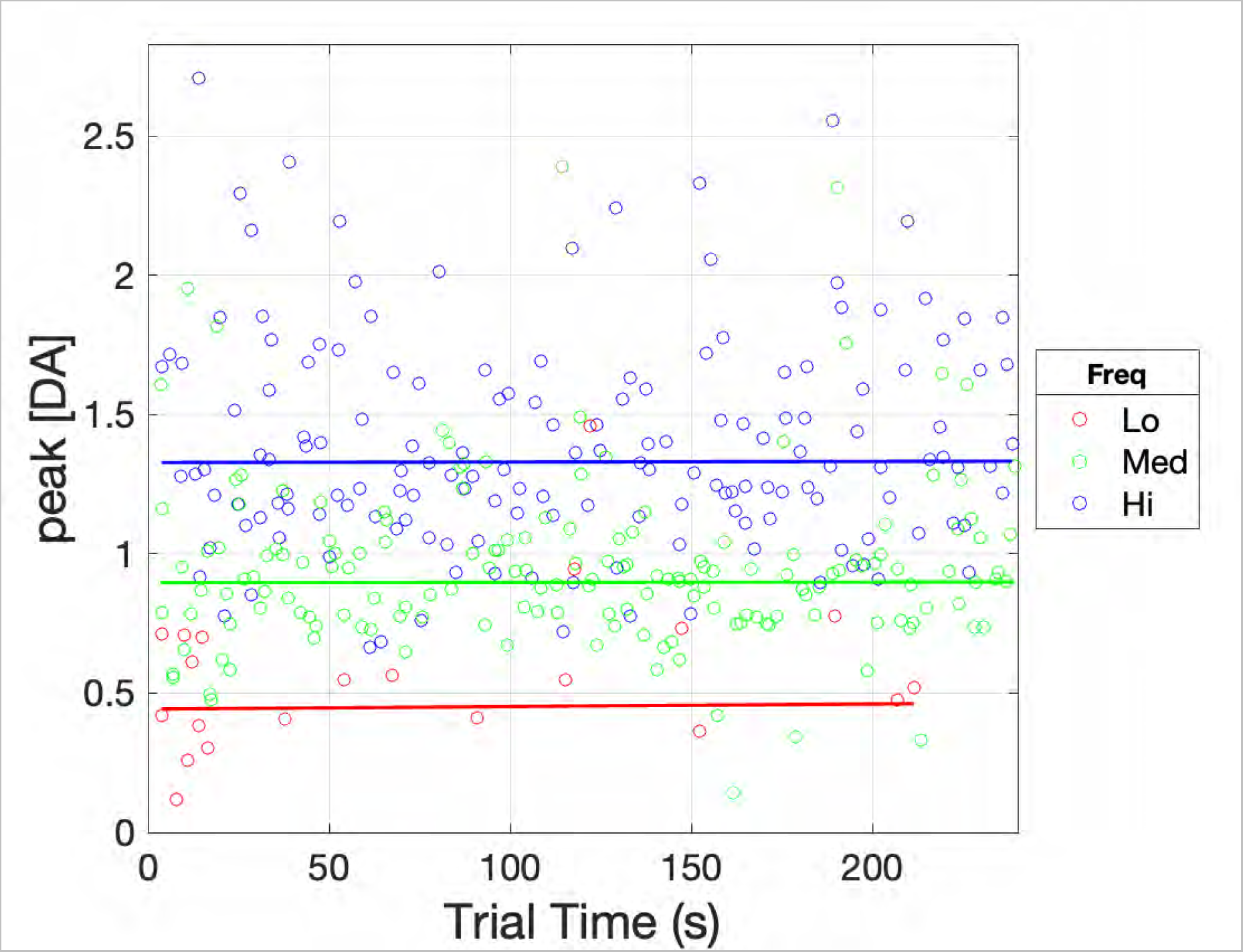
Peak dopamine concentration in response to each optical pulse train on test trials as a function of trial time. These are the same data that are shown as averages in Figures 2g and 2h. As in the case of the behavioral data in Figures 2e and 2f, multiple functions were fit to the data (straight lines, 2- and 3-segment piecewise-linear functions), and the best-fitting function was selected on the basis of the Akaike Information Criterion.

Video S1. A rat works for optical stimulation of its midbrain dopamine neurons. The black trace shows the estimated dopamine concentration at the surface of a carbon-fiber FSCV electrode in the nucleus accumbens terminal field. The blue trace shows each stimulation train as a single rectangle. Delivery of optical stimulation is indicated by the blue beam emanating from the laser symbol. In order to trigger the stimulation, the rat must accumulate 2 s on a clock by depressing the lever. On its first attempt, the rat releases the lever after ∼1.7 s and thus must return and press again in order to meet the criterion. On subsequent bouts, the rat succeeds in meeting criterion with a single lever depression.

Link to video:

https://www.biorxiv.org/content/biorxiv/early/2022/08/11/2022.08.08.502043/DC2/embed/media-2.mp4?download=true

## References

1. Montague, P. R., Dayan, P., Person, C. & Sejnowski, T. J. Bee foraging in uncertain environments using predictive hebbian learning. Nature 377, 725–728 (1995).

2. Montague, P. R., Dayan, P. & Sejnowski, T. J. A framework for mesencephalic dopamine systems based on predictive Hebbian learning. J Neurosci 16, 1936–1947 (1996).

3. Sutton, R. S. & Barto, A. G. Reinforcement learning: an introduction. (MIT Press, 2018).

4. Colombo, M. Deep and beautiful. The reward prediction error hypothesis of dopamine. Stud Hist Philos Biol Biomed Sci 45, 57–67 (2014).

5. Sutton, R. S. Learning to predict by the methods of temporal differences. Mach Learn 3, 9–44 (1988).

6. Waelti, P., Dickinson, A. & Schultz, W. Dopamine responses comply with basic assumptions of formal learning theory. Nature 412, 43–48 (2001).

7. Patriarchi, T. et al. Ultrafast neuronal imaging of dopamine dynamics with designed genetically encoded sensors. Science 360, eaat4422 (2018).

8. Sun, F. et al. A Genetically Encoded Fluorescent Sensor Enables Rapid and Specific Detection of Dopamine in Flies, Fish, and Mice. Cell 174, 481–496.e19 (2018).

9. Berridge, K. C. The debate over dopamine’s role in reward: the case for incentive salience. Psychopharmacology 191, 391–431 (2007).

10. Ludvig, E. A., Bellemare, M. G. & Pearson, K. G. A primer on reinforcement learning in the brain: Psychological, computational, and neural perspectives. in *Computational neuroscience for advancing artificial intelligence: Models*, methods and applications 111–144 (IGI Global).

11. Domesick, V. B., Stinus, L. & Paskevich, P. A. The cytology of dopaminergic and nondopaminergic neurons in the substantia nigra and ventral tegmental area of the rat: A light- and electron-microscopic study. Neuroscience 8, 743–765 (1983).

12. Yizhar, O., Fenno, L. E., Davidson, T. J., Mogri, M. & Deisseroth, K. Optogenetics in neural systems. Neuron 71, 9–34 (2011).

13. Redish, A. D. Addiction as a computational process gone awry. Science 306, 1944–1947 (2004).

14. Schultz, W., Dayan, P. & Montague, P. R. A Neural Substrate of Prediction and Reward. Science 275, 1593–1599 (1997).

15. Dolan, R. J. & Dayan, P. Goals and habits in the brain. Neuron 80, 312–325 (2013).

16. Daw, N. D., Gershman, S. J., Seymour, B., Dayan, P. & Dolan, R. J. Model-Based Influences on Humans’ Choices and Striatal Prediction Errors. Neuron 69, 1204–1215 (2011).

17. Sharpe, M. J. et al. Dopamine transients are sufficient and necessary for acquisition of model-based associations. Nat Neurosci 20, 735–742 (2017).

18. Wunderlich, K., Smittenaar, P. & Dolan, R. J. Dopamine Enhances Model-Based over Model-Free Choice Behavior. Neuron 75, 418–424 (2012).

19. Robinson, D. L., Hermans, A., Seipel, A. T. & Wightman, R. M. Monitoring rapid chemical communication in the brain. Chem Rev 108, 2554–2584 (2008).

20. Breton, Yannick-André. Molar and Molecular Models of Performance for Rewarding Brain Stimulation. (Concordia University, 2013).

21. Ahilan, S. et al. Learning to use past evidence in a sophisticated world model. PLOS Computational Biology 15, e1007093 (2019).

22. Pallikaras, V., Carter, F., Velazquez-Martinez, D. N., Arvanitogiannis, A. & Shizgal, P. The trade-off between pulse duration and power in optical excitation of midbrain dopamine neurons approximates Bloch’s law. Behavioural Brain Research 419, 113702 (2022).

23. Cartoni, E., Balleine, B. & Baldassarre, G. Appetitive Pavlovian-instrumental Transfer: A review. Neuroscience & Biobehavioral Reviews 71, 829–848 (2016).

24. Cover, C. G. et al. Whole brain dynamics during optogenetic self-stimulation of the medial prefrontal cortex in mice. Commun Biol 4, 66 (2021).

25. Hyland, B. I., Reynolds, J. N. J., Hay, J., Perk, C. G. & Miller, R. Firing modes of midbrain dopamine cells in the freely moving rat. Neurosci 114, 475–492 (2002).

26. Paladini, C. A. & Roeper, J. Generating bursts (and pauses) in the dopamine midbrain neurons. Neuroscience 282, 109–121 (2014).

27. Kuznetsova, A. Y., Huertas, M. A., Kuznetsov, A. S., Paladini, C. A. & Canavier, C. C. Regulation of firing frequency in a computational model of a midbrain dopaminergic neuron. J Comput Neurosci 28, 389–403 (2010).

28. Hollon, N. G. et al. Nigrostriatal dopamine signals sequence-specific action-outcome prediction errors. Current Biology (2021) doi:10.1016/j.cub.2021.09.040.

29. Gallistel, C., Shizgal, P. & Yeomans, J. A portrait of the substrate for self-stimulation. Psychol Rev 88, 228–273 (1981).

30. Trujillo-Pisanty, I., Conover, K., Solis, P., Palacios, D. & Shizgal, P. Dopamine neurons do not constitute an obligatory stage in the final common path for the evaluation and pursuit of brain stimulation reward. PLOS ONE 15, e0226722 (2020).

31. Gallistel, C. R. & Gibbon, J. Time, rate, and conditioning. Psychological Review 107, 289–344 (2000).

32. Gallistel, C. R., Craig, A. R. & Shahan, T. A. Contingency, contiguity, and causality in conditioning: Applying information theory and Weber’s Law to the assignment of credit problem. Psychological Review 126, 761–773 (2019).

33. Gallistel, C. R. & Latham, P. E. Bringing Bayes and Shannon to the Study of Behavioural and Neurobiological Timing and Associative Learning. Timing Time Percept. 11, 29–89 (2022).

34. Balsam, P. D. & Gallistel, C. R. Temporal maps and informativeness in associative learning. Trends in Neurosciences 32, 73–78 (2009).

35. Hernandez, G., Breton, Y.-A., Conover, K. & Shizgal, P. At what stage of neural processing does cocaine act to boost pursuit of rewards? PLoS ONE 5, (2010).

36. Solomon, R. B., Conover, K. & Shizgal, P. Valuation of opportunity costs by rats working for rewarding electrical brain stimulation. PLOS ONE 12, e0182120 (2017).

37. Namboodiri, V. M. K. How do real animals account for the passage of time during associative learning? Behavioral Neuroscience 136, 383–391 (2022).

38. Breton, Y.-A., Mullett, A., Conover, K. & Shizgal, P. Validation and extension of the reward-mountain model. Front Behav Neurosci 7, 125 (2013).

39. Arvanitogiannis, A. & Shizgal, P. The reinforcement mountain: allocation of behavior as a function of the rate and intensity of rewarding brain stimulation. Behav Neurosci 122, 1126– 1138 (2008).

40. Simmons, J. M. & Gallistel, C. R. Saturation of subjective reward magnitude as a function of current and pulse frequency. Behav Neurosci 108, 151–160 (1994).

41. Wang, J. X. et al. Prefrontal cortex as a meta-reinforcement learning system. Nature Neuroscience (2018) doi:10.1038/s41593-018-0147-8.

42. Jeong, H. et al. Mesolimbic dopamine release conveys causal associations. Science 378, eabq6740 (2022).

43. Burke, D. A. et al. Few-shot learning: temporal scaling in behavioral and dopaminergic learning. 2023.03.31.535173 Preprint at 10.1101/2023.03.31.535173 (2023).

44. Garr, E. et al. Mesostriatal dopamine is sensitive to specific cue-reward contingencies. 2023.06.05.543690 Preprint at 10.1101/2023.06.05.543690 (2023).

45. Blanco-Pozo, M., Akam, T. & Walton, M. Dopamine reports reward prediction errors, but does not update policy, during inference-guided choice. bioRxiv 2021.06.25.449995 (2021) doi:10.1101/2021.06.25.449995.

46. Trujillo-Pisanty, I., Sanio, C., Chaudhri, N. & Shizgal, P. Robust optical fiber patch-cords for in vivo optogenetic experiments in rats. MethodsX 2, 263–271 (2015).

47. Breton, Y.-A., Marcus, J. C. & Shizgal, P. Rattus Psychologicus: construction of preferences by self-stimulating rats. Behav Brain Res 202, 77–91 (2009).

48. Trujillo-Pisanty, I., Solis, P., Conover, K., Dayan, P. & Shizgal, P. On the forms of learning supported by rewarding optical stimulation of dopamine neurons. in Society for Neuroscience Abstract Viewer 66.06 (2016).

49. Akaike, H. A new look at the statistical model identification. IEEE Transactions on Automatic Control 19, 716–723 (1974).

50. Rodeberg, N. T. et al. Construction of Training Sets for Valid Calibration of in Vivo Cyclic Voltammetric Data by Principal Component Analysis. Anal. Chem. 87, 11484–11491 (2015).

51. Kishida, K. T. et al. Subsecond dopamine fluctuations in human striatum encode superposed error signals about actual and counterfactual reward. PNAS 113, 200–205 (2016).

52. Keithley, R. B. & Wightman, R. M. Assessing Principal Component Regression Prediction of Neurochemicals Detected with Fast-Scan Cyclic Voltammetry. ACS Chem Neurosci 2, 514–525 (2011).

53. Heien, M. L. A. V., Johnson, M. A. & Wightman, R. M. Resolving neurotransmitters detected by fast-scan cyclic voltammetry. Anal Chem 76, 5697–5704 (2004).

54. Cossette, M.-P. Anatomical and computational models of the role of phasic dopamine signaling in intracranial self-stimulation: psychophysical and electrochemical tests. (Concordia University, 2019).

## References

1. Pallikaras, V., Carter, F., Velazquez-Martinez, D. N., Arvanitogiannis, A. & Shizgal, P. The trade-off between pulse duration and power in optical excitation of midbrain dopamine neurons approximates Bloch’s law. Behavioural Brain Research 419, 113702 (2022).

