## Supporting information for "Does phasic dopamine release cause policy updates?"

[1] Department of Psychology, Concordia University, Montreal, QC, Canada
[2] Montreal Institute for Learning Algorithms, Université de Montréal, Montreal, QC, Canada
[3] Department of Psychology, Langara College, Vancouver, BC, Canada

[&] corresponding author

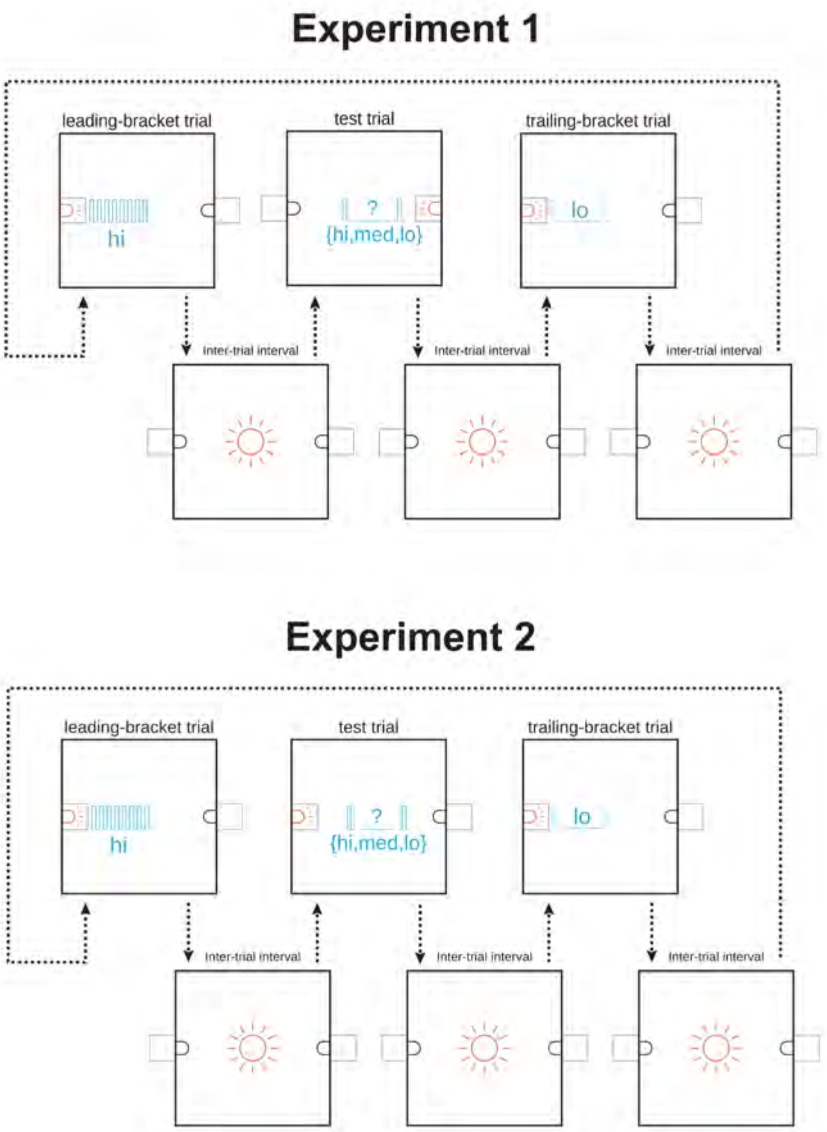


Figure S1: Testing paradigms. In Experiment 1, different levers were used to trigger stimulation delivery on bracket (upper left and right panels) and test (upper middle panel) trials. Stimulation strengths (pulse frequencies) are shown in cyan. The paradigm employed in Experiment 2 is identical to the one employed in Experiment 1 with one exception: the same lever was used to trigger stimulation delivery on bracket (upper left and right panels) and test (upper middle panel) trials.

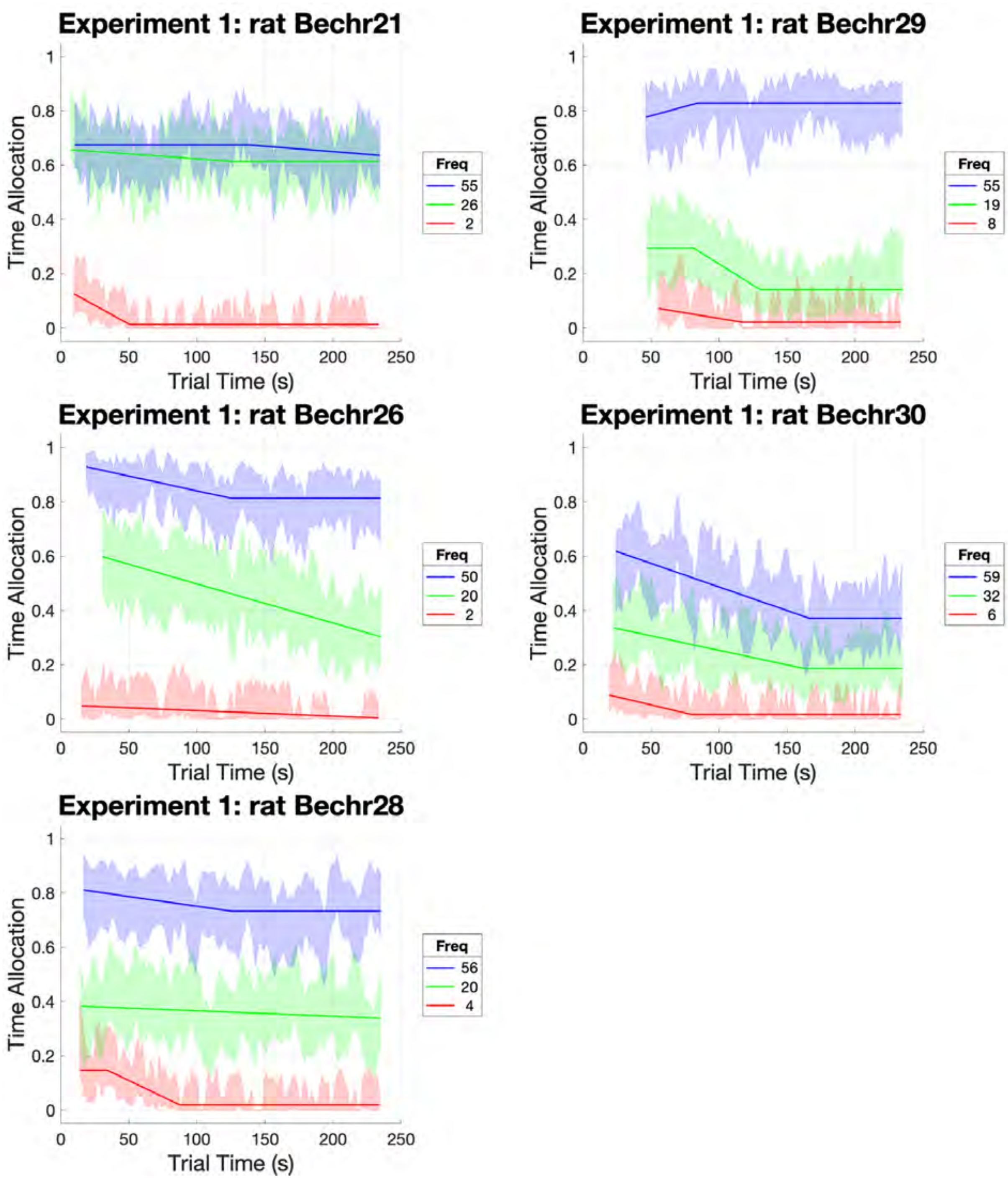


Figure S2: Stability of test-trial responding for rewarding optical stimulation for each subject in Experiment 1. The proportion of time that the lever was depressed is shown in successive 2-s time blocks. Lines show best-fitting piecewise linear functions, and error bands show the surrounding 95% confidence intervals. The stimulation strength was set by the pulse frequency, which was either high, medium, or low. The plots for each stimulation strength begin at the time block when responding had commenced on 75% of the trials and end at the 3rd to last time block.

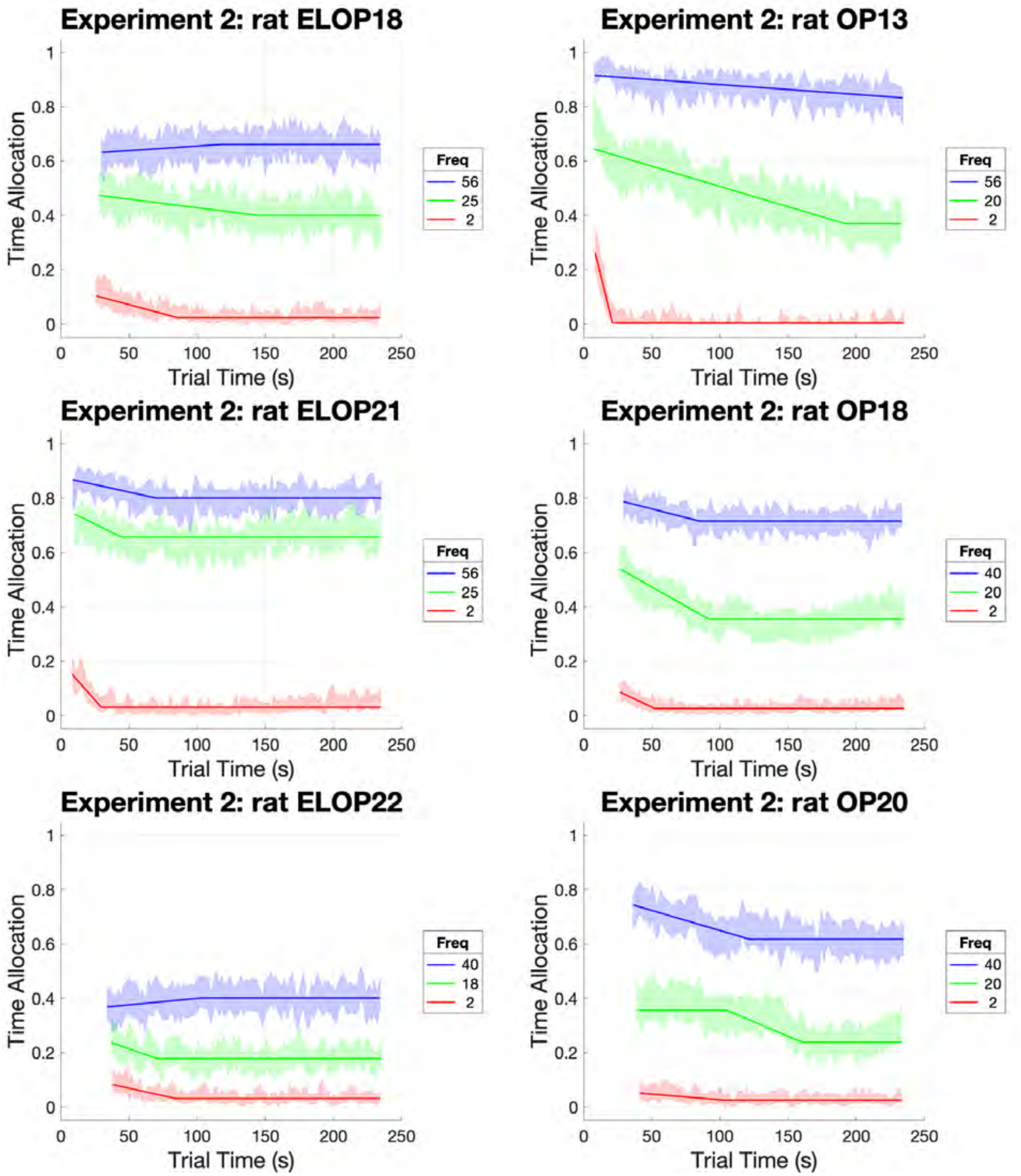


Figure S3: Stability of test-trial responding for rewarding optical stimulation for each subject in Experiment 2. For details, see the caption for Figure S2.

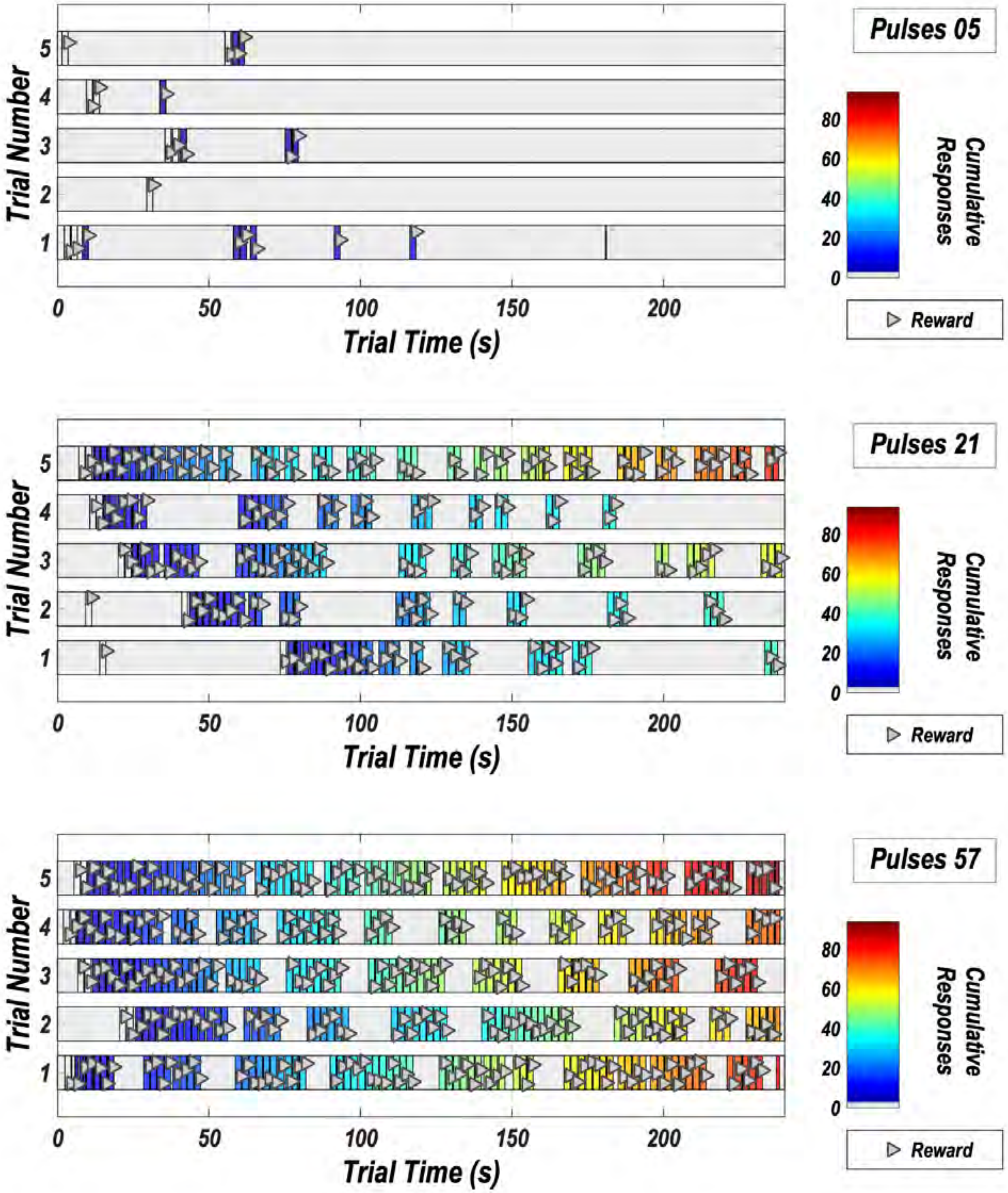


Figure S4: Pattern of responding in Experiment 1. The data are from rat Bechr28, test session 3. Responding for the low, medium and high stimulation strength is shown in the upper, middle, and bottom panels, respectively. Bout durations, and response totals (color map) typically increase as a function of increasing stimulation strength, whereas inter-bout pauses typically decrease. The triangles indicate lever presses and are jittered vertically to increase visibility.

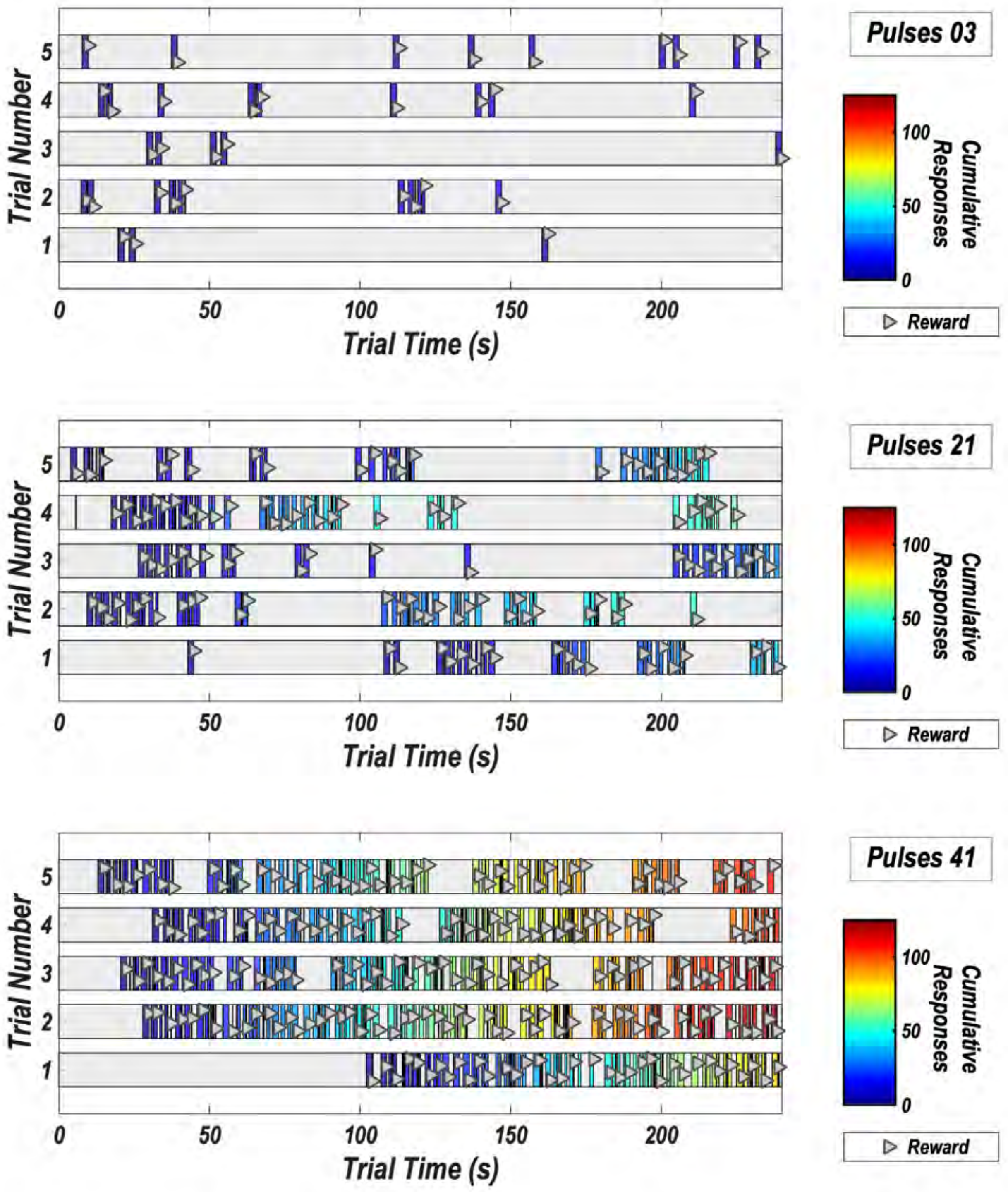


Figure S5: Pattern of responding in Experiment 2. The data are from rat OP18, test session 5. Responding for the low, medium and high stimulation strength is shown in the upper, middle, and bottom panels, respectively. Bout durations, and response totals (color map) typically increase as a function of increasing stimulation strength, whereas inter-bout pauses typically decrease. The triangles indicate lever presses and are jittered vertically to increase visibility.

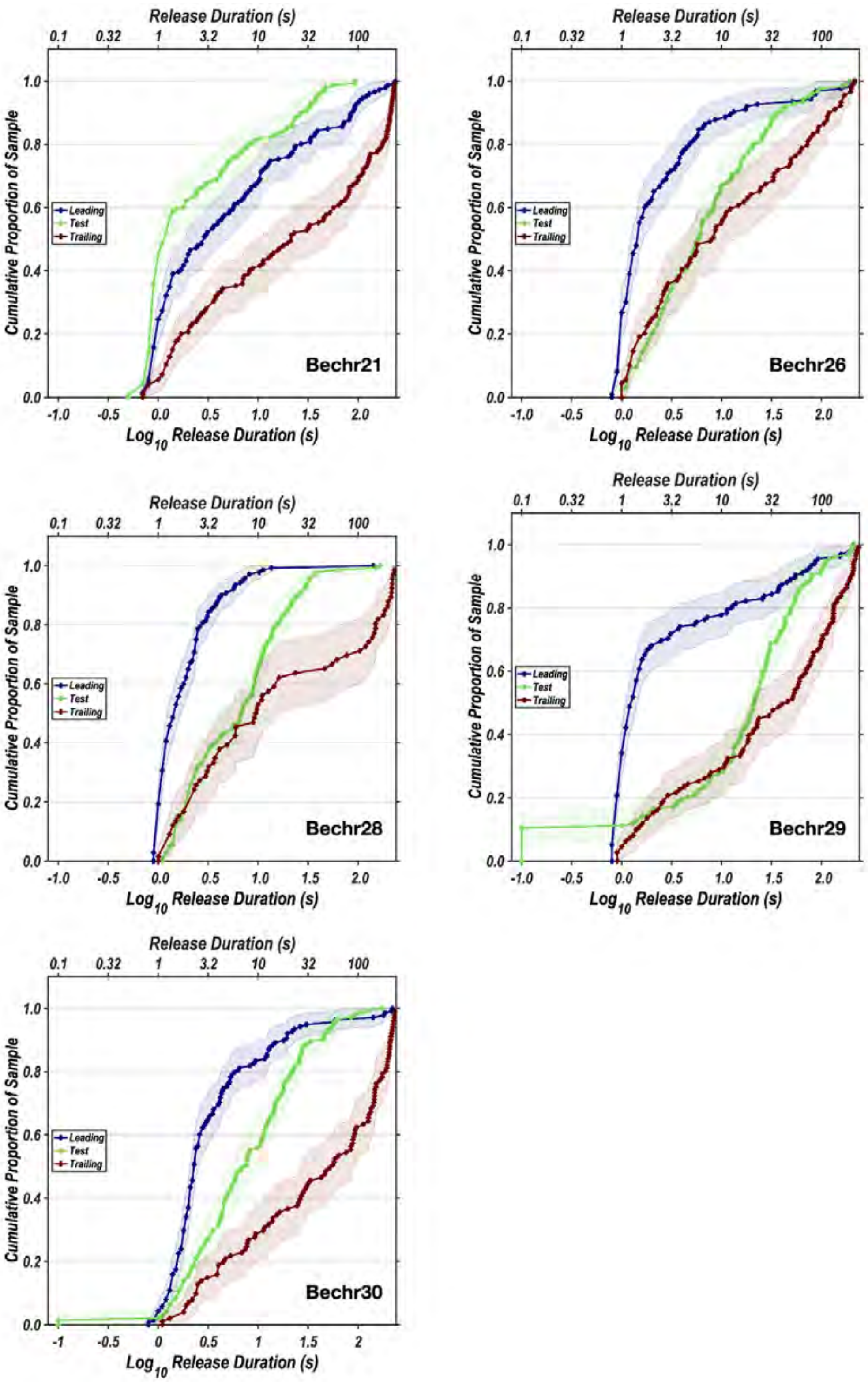


Figure S6: Cumulative distributions of the initial latencies on all trial types in Experiment 1.

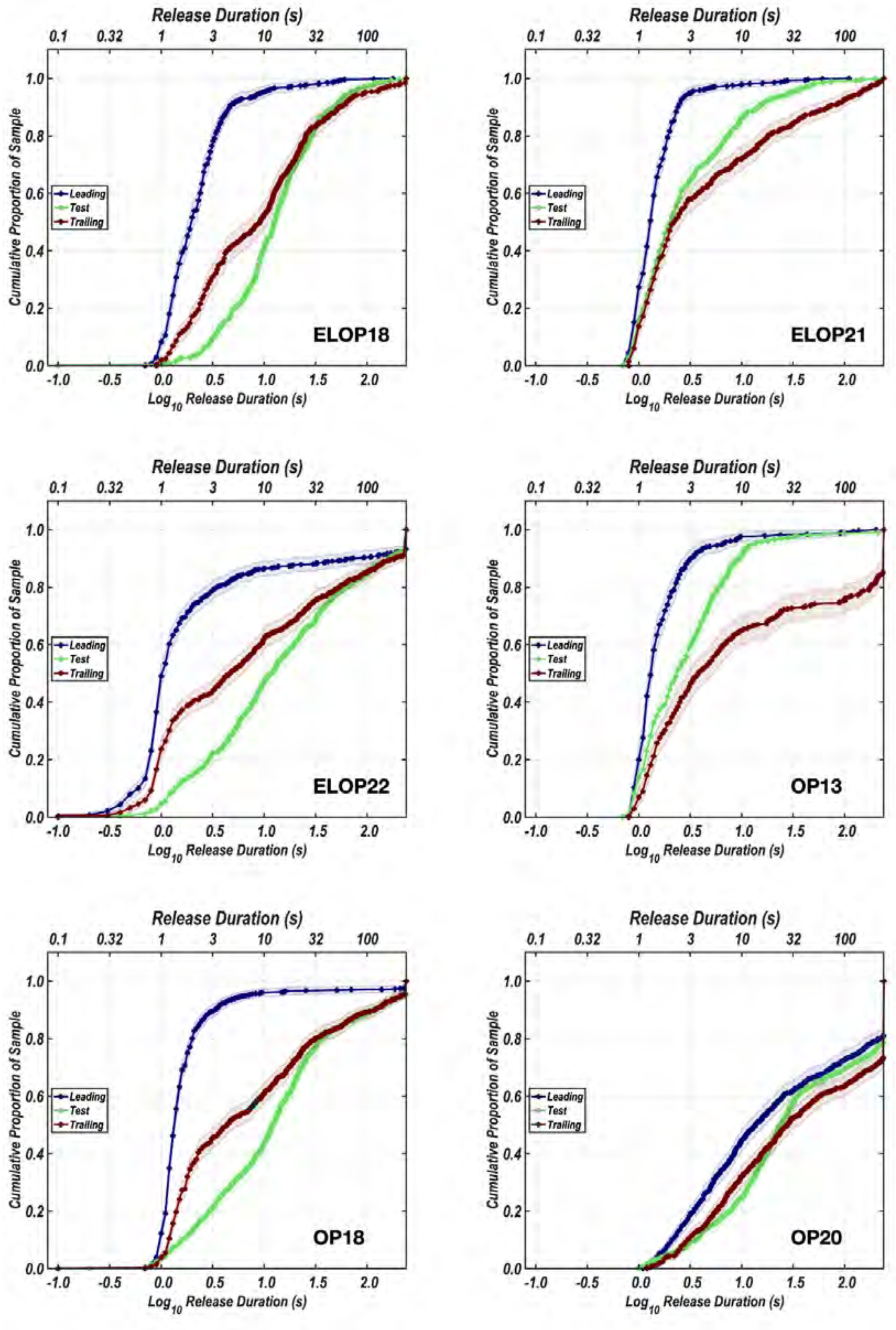


Figure S7: Cumulative distributions of the initial latencies on all trial types in Experiment 2.

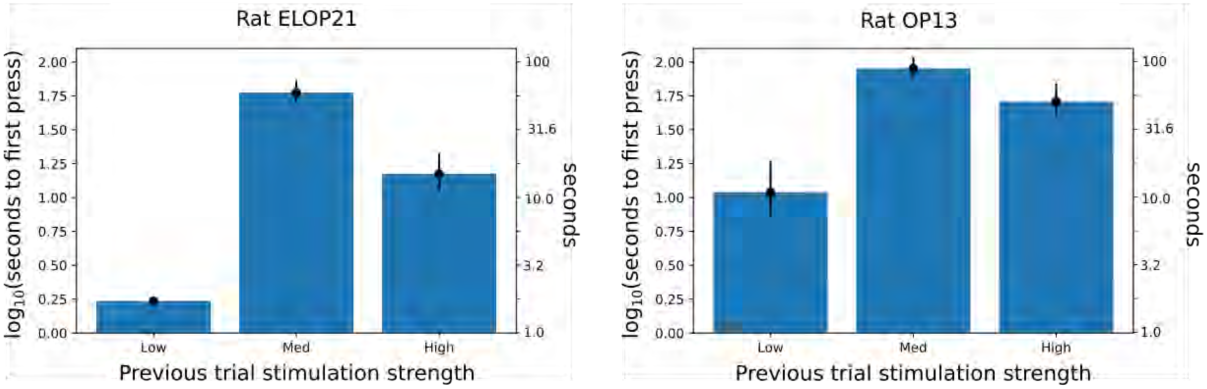


Figure S8: Bar plot of latencies until first press on trailing trials depending on the stimulation strength on the previous test trial. Data are from the two rats that learned the entire triadic trial structure.

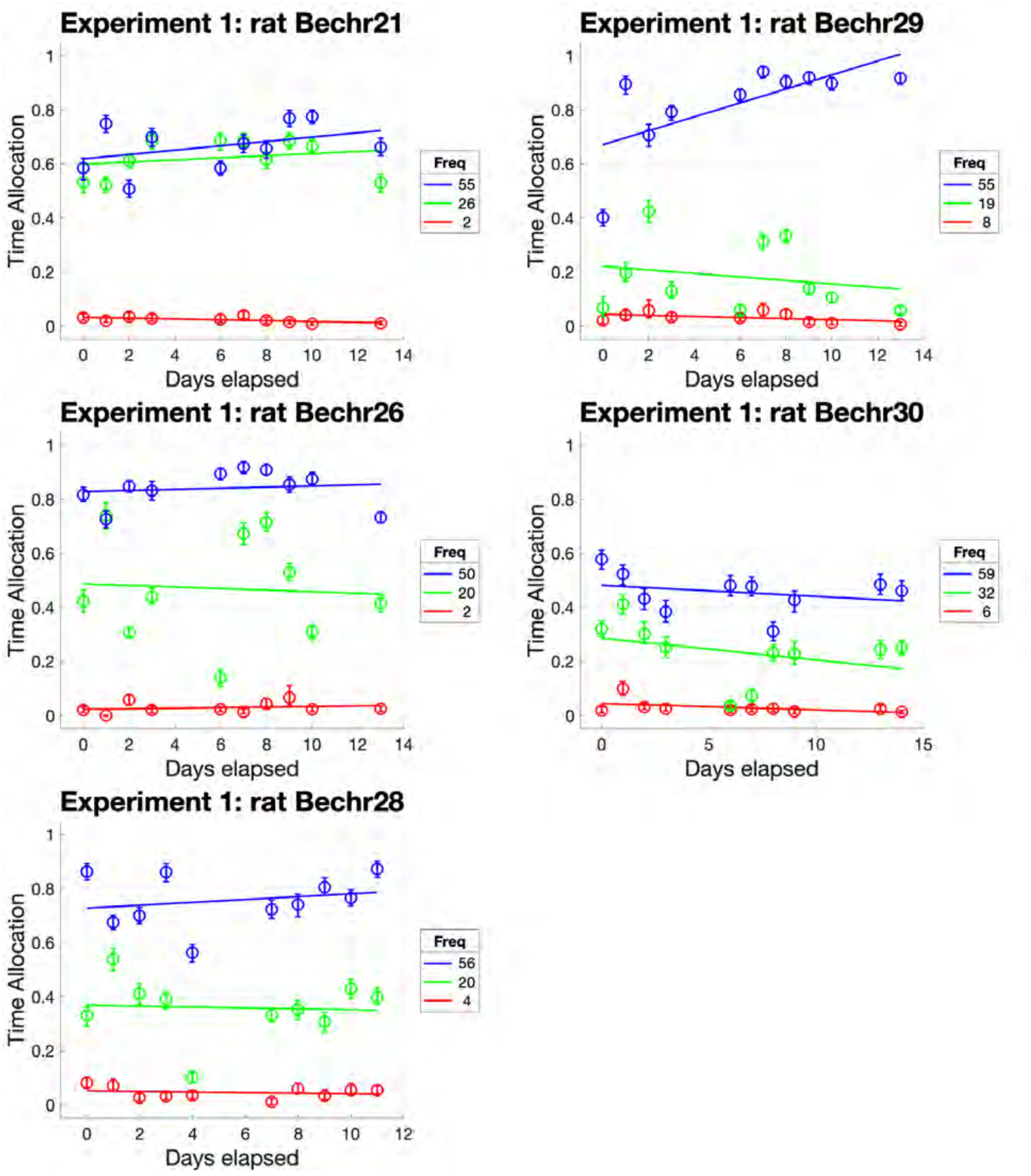


Figure S9: Evolution of time spent working as a function of stimulation strength over the course of Experiment 1. Many more sessions were run in Experiment 2, thus extending the period over which drift could be observed.

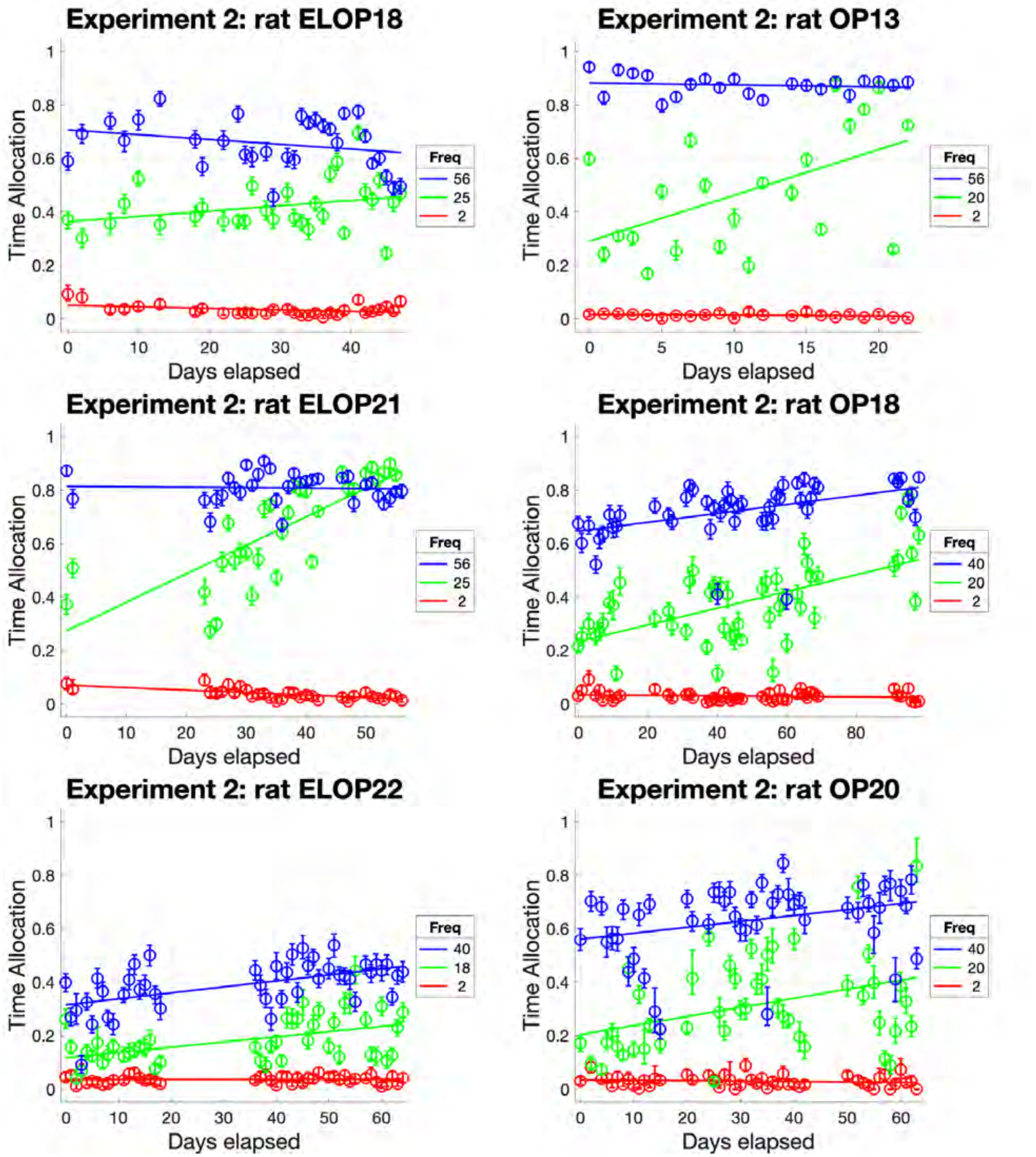


Figure S10: Evolution of time spent working as a function of stimulation strength over the course of Experiment 2. Over the prolonged period of repeated testing, time allocated to working for the medium-strength reward increased reliably in all rats, save for ELOP18. Data points are means over the five trials carried out at each stimulation strength during each test session.

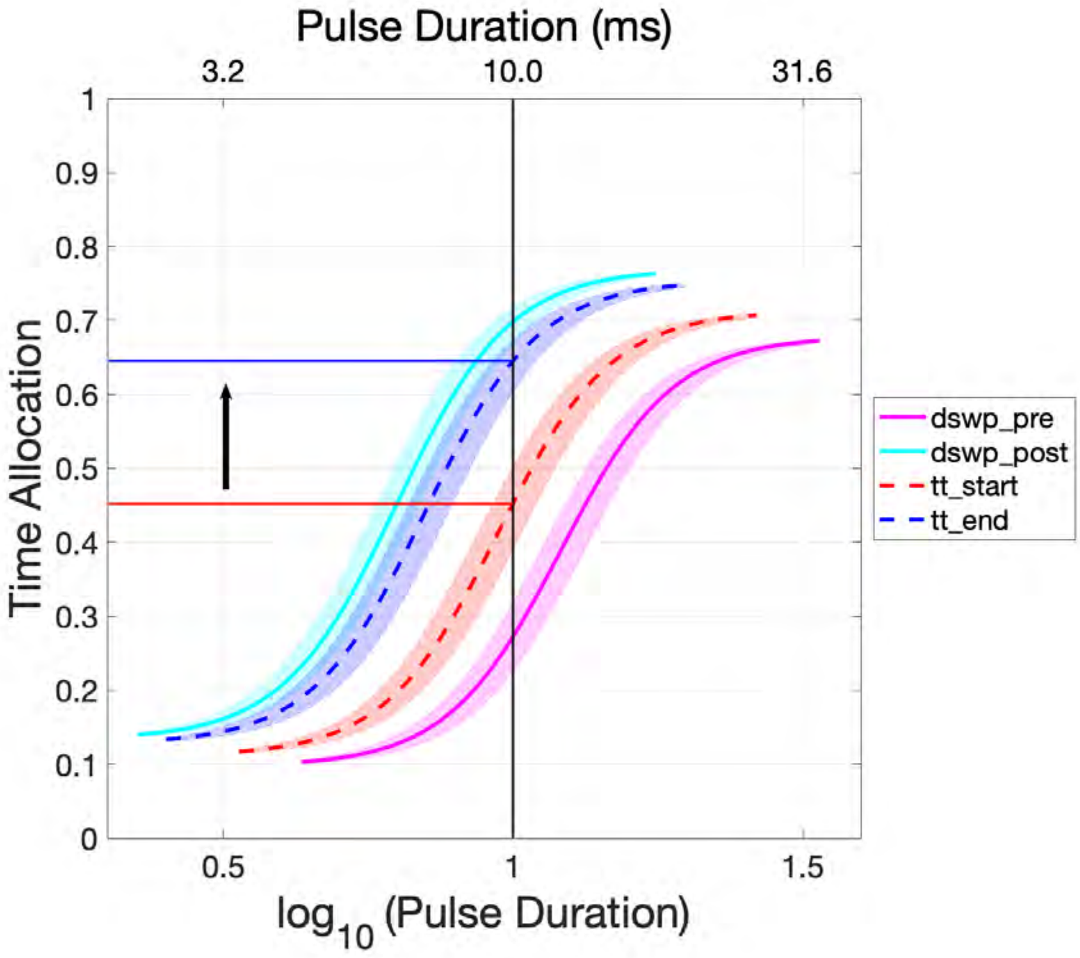


Figure S11. Fitted and interpolated sigmoidal functions. The solid curves represent sigmoidal functions fitted to pulse-duration sweeps obtained from rat OP18 prior (dswp-pre) and after (dswp-post) triadic-trial testing. The dashed curves are interpolated sigmoids for the start (tt-start) and end (tt-end) of triadic-trial testing. These were generated on the basis of the slope of drift in Figure 4b, the start- and end- dates of triadic-trial testing, and the parameters of the sigmoids fitted to the data from the pre- and post- triadic-trial pulse-duration sweeps. Shaded areas denote the 95% confidence intervals surrounding the position-parameter estimates. The solid blue and red lines show the predicted change in time allocation over the course of triadic trial testing, which was carried out at a pulse duration of 10 ms.

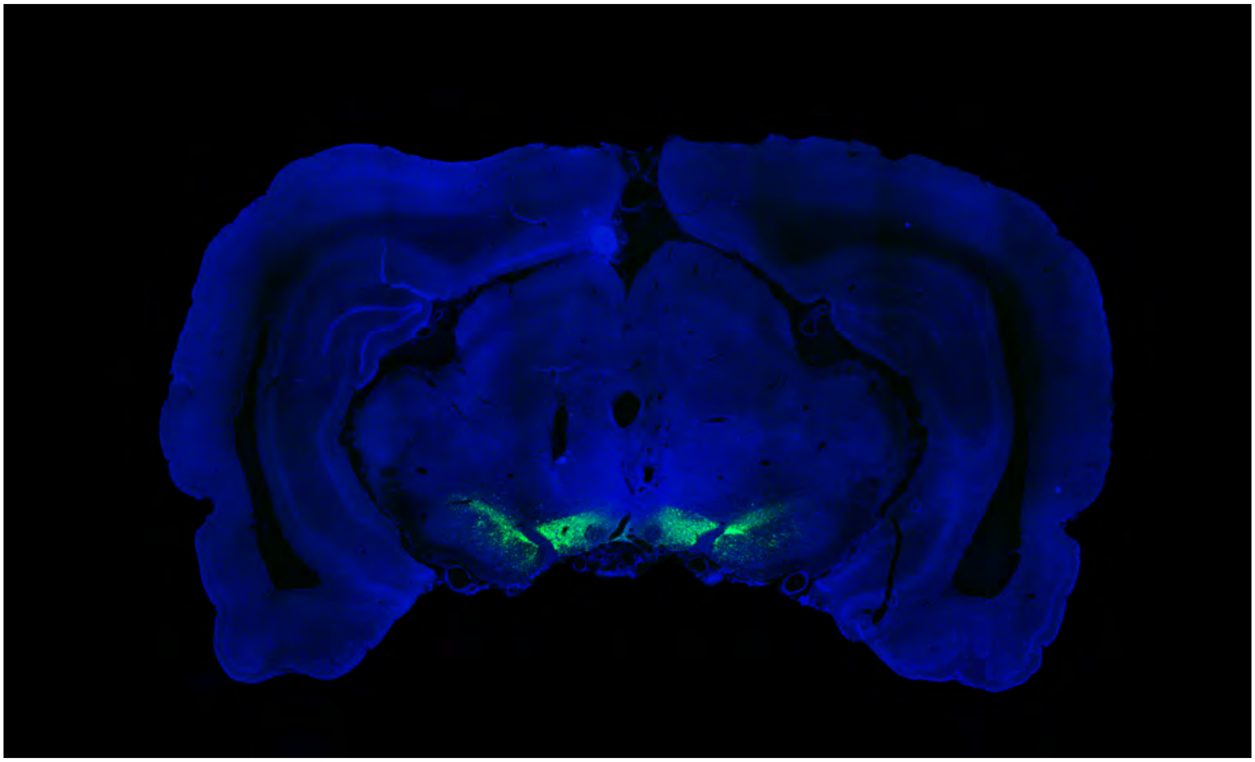

Figure S12. Histological image for subject OP13 (Experiment 2); eYFP expression (green) shown along with DAPI (blue) for anatomical reference (image enhanced by rescaling the distribution of pixel intensities). This figure was published previously[1].

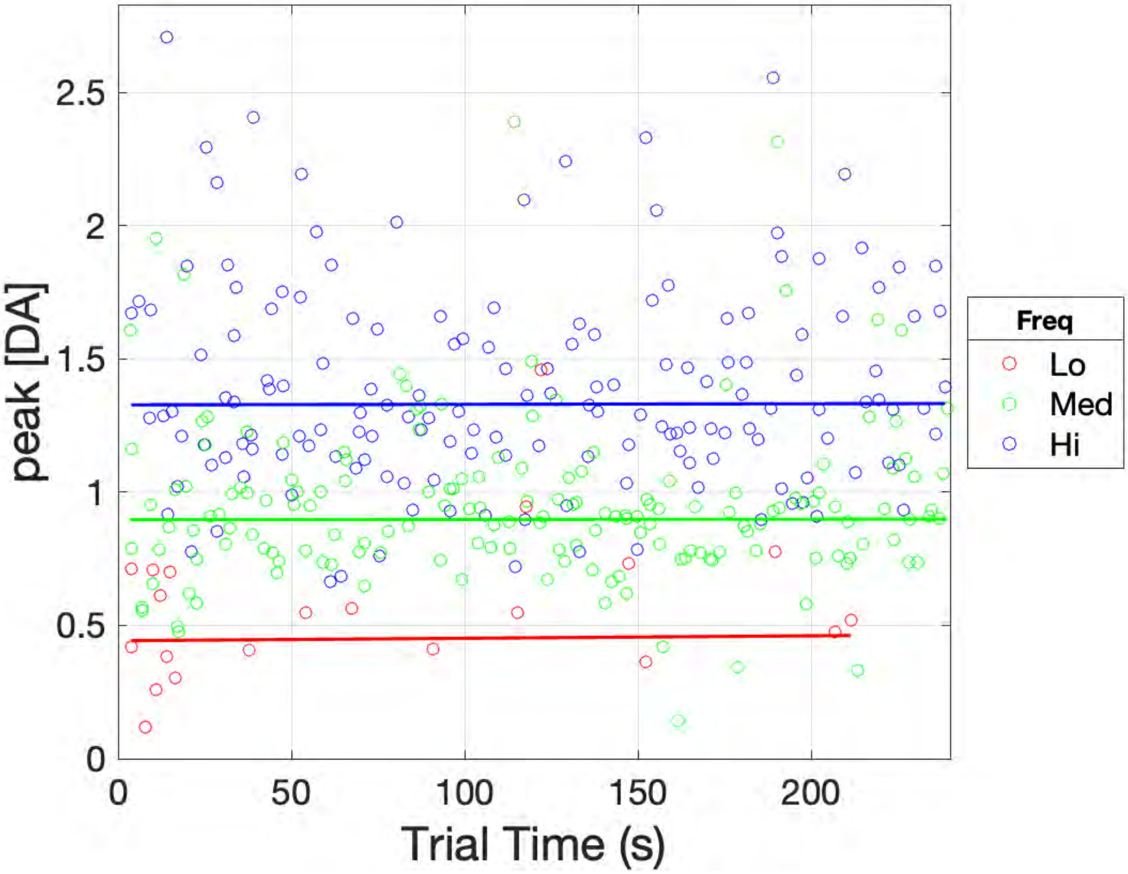


Figure S13. Peak dopamine concentration in response to each optical pulse train on test trials as a function of trial time. These are the same data that are shown as averages in Figures 2g and 2h. As in the case of the behavioral data in Figures 2e and 2f, multiple functions were fit to the data (straight lines, 2- and 3-segment piecewise-linear functions), and the best-fitting function was selected on the basis of the Akaike Information Criterion.

Video S1. A rat works for optical stimulation of its midbrain dopamine neurons. The black trace shows the estimated dopamine concentration at the surface of a carbon-fiber FSCV electrode in the nucleus accumbens terminal field. The blue trace shows each stimulation train as a single rectangle. Delivery of optical stimulation is indicated by the blue beam emanating from the laser symbol. In order to trigger the stimulation, the rat must accumulate 2 s on a clock by depressing the lever. On its first attempt, the rat releases the lever after ~1.7 s and thus must return and press again in order to meet the criterion. On subsequent bouts, the rat succeeds in meeting criterion with a single lever depression.

Link to video:
https://www.biorxiv.org/content/biorxiv/early/2022/08/11/2022.08.08.502043/DC2/embed/media-2.mp4?download=true

### References

1. Pallikaras, V., Carter, F., Velazquez-Martinez, D. N., Arvanitogiannis, A. & Shizgal, P. The trade-off between pulse duration and power in optical excitation of midbrain dopamine neurons approximates Bloch's law. *Behavioural Brain Research* **419**, 113702 (2022).
